# Evidence of spatial regime shifts: contrasting spatial dynamics of three Atlantic cod (*Gadus* morhua) populations in the North Sea

**DOI:** 10.1101/2023.10.13.562168

**Authors:** Enrico Cecapolli, Giovanni Romagnoni, Alexandra M. Blöcker, Christian Möllmann, Camilla Sguotti

## Abstract

Recent scientific assessments have started to consider the presence of multiple populations of Atlantic cod (*Gadus morhua*) in the North Sea due to genetic evidence. The Atlantic cod stock as a whole has experienced a regime shift in the North Sea and is now considered to be in a persistent depleted state. Here we investigated if the regime shift detected for North Sea cod manifested differently across areas (South, North-West and Viking (North-East)), revealing different dynamics and recovery potential for the corresponding populations. Methods from change point analyses to stochastic cusp models were applied to detect the regime shifts. We found that cod in the Southern North Sea has undergone a regime shift and is now in a persistent depleted state. The Viking population is in a low state but with high resilience, making its dynamics more predictable and thus easier to manage. Finally, the north-western population did not show signs of a regime shift and is in a high biomass level. Several abiotic and biotic drivers (e.g. fishing, warming) contribute to these differences. Our study provides useful information for the restructuring of management units of cod in the North Sea and highlights the importance of looking at regime shifts spatially.

## Introduction

Regime shifts, i.e. unexpected and often sudden changes between system states, are among the most debated dynamics of marine ecosystems [1–3]. Systems that undergo regime shifts change from a past to a new state, characterized by different ecological structures and functions and by a certain degree of persistence (i.e. hysteresis) over time [4–7]. Resilience, another fundamental property of systems that undergo regime shifts, is defined as the capability of a system to maintain its structure and functions even when impacted. As a general rule, systems that undergo regime shifts have a lower resilience. Regime shifts are still difficult to characterize because, following non-linear discontinuous dynamics, they are highly unpredictable and often not easily detectable using simple statistical models [8–10]. Two inherent limitations regarding regime shifts lie also in the usage of appropriate temporal and spatial (or hierarchical) scales when trying to detect them [11–14]. While the temporal scale of regime shifts is often considered, the spatial/hierarchical scale is generally neglected [15]. This limitation resides in the difficulty of identifying the study unit of the system and what its limits are. For instance, while it is rather easy to study a coral reef, if we study an ecosystem or a population it is always difficult to determine the spatial boundaries of the system. Indeed, a regime shift that is detected within a larger area could in fact have happened in a smaller portion of the area, or just involve specific habitats within the overall area or just a subpopulation. Therefore, within a spatial or hierarchical unit different states of the system could exist, with different hysteresis (persistence) levels [1,4]. The spatial context of regime shifts is extremely important to explore, especially when developing management measures [16–18]. Thus, a key for a good understanding of regime shifts should also include a punctual consideration of the proper spatial scale, to avoid misleading results.

The North Sea is a paradigmatic example of a large-scale ecosystem that underwent a regime shift encompassing most of its trophic web from plankton to large predatory fishes [19–23]. The regime shift occurred in particular due to the increase of temperature in the 1980s and the changes in hydro-climatic forces, coupled with unsustainable fishing pressure especially on some target fish species [20,24]. The regime shift started at the bottom of the trophic web with changes in composition and abundances of the phytoplankton and zooplankton communities, and then cascade to higher trophic-level organisms such as economically important fishes [19,20,25,26]. Among the North Sea fish species, Atlantic cod (*Gadus morhua*), one of the most iconic species in the world, experienced a strong depletion due to overexploitation, coupled with the disappearance of its main prey zooplankton species [26,27]. Regime shifts analyses conducted on the North Sea cod stock have demonstrated that the stock experienced a regime shift due to fishing and climate change and is at present in a persistent depleted state characterized by low resilience [28,29].

The regime shift of Atlantic cod in the North Sea started around the 1980s, when the stock began to decline and reached a very low and depleted biomass level in the 2000 (ICES 2019). At present times, the stock shows limited recovery compared to previous abundances [29], still being below sustainable reference levels [30]. The recovery potential of Atlantic cod in the North Sea is low: changes in temperature due to climate change, and the reorganization of the full trophic assemblage contribute to the creation of hysteresis. However, in the northern areas of the North Sea, an increase of the population is occurring, most likely because of locally lower sea temperatures, while in the warmer southern areas the species seems to suffer [31]. These observations suggest that the consideration of a single stock unit for management in the North Sea might be inappropriate since populations exist in the Southern, North-Western, and Viking (North-Eastern) areas [32–35]. The previously detected regime shift encompassing the entire stock could have occurred differently among the spatial units considered, resulting in different recovery potential and hysteresis patterns, and thus rendering management measures applied to the full basin mostly ineffective. In this context, management units utilized for stock assessment have been reviewed and the most recent benchmark proposes a multi-stock assessment model application [36]. However, while the inclusion of stock structure allows specific dynamics to be captured, the individual stocks’ past exposition to irreversible regime shifts, their current proximity to thresholds and their resilience to multiple pressures, remain unexplained. Thus, a comprehensive spatial analysis of regime shifts in the putative North Sea cod populations is fundamental to understand how recovery could be enhanced, and what populations are more likely to respond positively to management measures.

Here we analyze whether different dynamics (linear, non-linear, discontinuous), and especially regime shifts dynamics (non-linear and discontinuous), occurred to three putative sub-populations of the cod stock: the Southern, North-Western, and Viking (North-Eastern) North Sea. This division has been defined based on genetic, ecological, and behavioral evidence (summarized in ICES WGNSSK report [30]). Firstly, we analyzed the populations’ development over time. Afterwards, we detected a number of possible drivers of the changes, and we tested for regime shifts dynamics using the stochastic cusp model, an approach deriving from catastrophe theory. The approach used in this investigation could result in useful hints for management purposes, as it gives a clearer overview of the status of the populations of cod and their dynamics over time in the spatial units considered, supporting the need to switch to a more effective strategy to support the recovery of the Atlantic Cod in the North Sea. Our study opens up new perspectives about spatial regime shifts and their importance.

## Material and Methods

### Data

Our study aims at understanding the dynamics of Atlantic cod (*Gadus morhua*) in three areas of the North Sea, which are believed to represent three distinct populations of the stock [34]. The three areas of focus are: the Southern, North-Western, Viking (North-Eastern) North Sea, and were defined accordingly to stock assessment reports [30,34]. All the analyses described below were conducted separately for the three areas in order to analyze the different populations. We also repeated the same analyses by aggregating together the Southern and the North-Western area since it is still debated whether they can be treated together or separated due to their similarities in genetic traits and habitats [30,34,35] (See Supporting Information).

#### Cod population and community data

The estimates of cod abundance and the characterization of the fish community in the different areas were obtained using survey data coming from the North Sea International Bottom Trawl Survey (NS-IBTS), available from the International Council for the Exploration of the Sea (ICES; DATRAS, http://www.ices.dk/data/data-portals/Pages/DATRAS.aspx). Such survey is carried out twice a year (1st and 3rd quarters) since 1997 by the ICES, but started already with at least one survey in quarter 1 from the 1960’s, performed using a standard otter trawl as sampling gear for each vessel involved. Due to consistent sampling, we extracted the 1st quarter data (Q1, from January to March). Since the main focus of our analysis was Atlantic Cod, we first created a dataset of Catch per Unit of Effort (CPUE) of cod per area per year, and then a dataset of the total community per area per year that we used as the possible driver of changes in the cod populations. We used CPUE values of more than 100 species of the North Sea basin. In order to obtain a value of CPUE per area per year, we first summed specimens for every species in every station. Then we averaged the CPUE per ICES statistical rectangle (which are ICES geographical units) and finally we summed the values per area. In addition, to perform a more robust characterization of the community, we removed all species that presented a high number of non-valid values (NAs ≥ 15), and we restricted the dataset from 1985 to 2015 (years from which we had data for all the drivers).

#### Biotic variables

As biotic factors possibly related to cod dynamics, we used the quantity of big copepods, which included mainly *Calanus finmarchicus* and *Calanus helgolandicus*, since they are fundamental prey items for cod larvae[37]. Big copepods data came from the Continuous Plankton Recording (CPR), supplied by the Continuous Plankton Recorder program (www.cprsurvey.org) [38]. The values were provided by ICES rectangles, thus we averaged them to obtain estimates of their quantity per area per year.

#### Environmental drivers

As abiotic factors we used three environmental indices which are well known to impact fish populations: Sea Surface Temperature (SST), that was used as a proxy of climate change, Sea Bottom Temperature (SBT), which is most influential for adult cod since it is a demersal species, and Sea Surface Salinity (SSS), that can have an impact on a variety of fish species, even if direct effects on cod are not extensively reported. Data were available per rectangle, and we averaged all the values to obtain one estimate per area per year. These data were produced by the oceanographic model by Núñez-Riboni & Akimova [39].

#### Fishing effort

Finally, we checked whether different patterns of fishing effort could influence the dynamics of cod. Fishing effort data were taken from Couce et al. (2019) and covered the 31-year series from 1985 to 2015. These data are estimates of the hours fished by beam and otter trawler per ICES rectangle in the North Sea and are therefore a good estimate of spatial fishing effort [40]. In order to obtain yearly estimates per area we summed the hours trawled every year per area.

### Methods

All analyses were performed in the three areas (Southern, North-Western and Viking). We applied a range of analyses in order to be able to detect the presence of discontinuous dynamics in cod’s populations, along the lines of the protocol outlined in Sguotti et al. (2022) for the multi-method detection of regime shifts in marine systems.

#### Change point analysis

At first, we looked at the dynamics of Atlantic cod in the different areas by plotting its CPUE per year. We subsequently analyzed the cod time series to identify any proxy of the presence of abrupt changes, using three different types of change point analyses: the Bayesian change point analysis, the change in mean, and the change in both mean and variance [8,10,41]. These techniques allow to detect statistically relevant changes in the time series by computing a moving average (or variance) of its values and then checking when there is a significant difference. The change points in the time series were identified by combining the three methods and provided a first indication of a possible presence of regime shift.

#### Community analysis

To compare the trends of cod to the trend of the fish community, we performed a Principal Component Analysis (PCA), a multivariate technique to simplify complex datasets extracting their main variability [42]. In this case, as representative of the community we used the first two components, namely PC1 and PC2, which account for the majority of the variability [42]. To detect the presence of different community stages, we conducted a Cluster Analysis [43]. This classification technique allows to group data and find possible discontinuities in a dataset. For our purposes, we used resemblance matrices calculated with the Euclidean distance. The Cluster Analysis was based on the Ward’s criterion and the graphical interpretation of the obtained dendrogram allowed the construction of the final biplot with the best subdivision of the years. PC1 and PC2 loadings were extracted to obtain a simplified time series of the community in a unique plot.

#### Detecting hysteresis, driver-state plot

As a second step to investigate the presence of regime shift dynamics in the three cod populations, we graphically investigated the driver-state plots [8,15]. If a linear relationship was found, i.e. for every level of the driver there was just one level of the state variable, then the dynamic of the population was probably linear. Instead, if multiple levels of the state variable were present for one level of the driver, there could be an indication of discontinuous dynamics and thus a regime-shift-like behavior [10]. We plotted as the state variable CPUE of cod in the three areas and as the driver fishing effort, being the most relevant driver responsible for the decrease of the populations[29,31].

#### The stochastic cusp model

Finally, to prove the presence of discontinuous dynamics we performed stochastic cusp models for each area [44]. The stochastic cusp model is derived from the catastrophe theory, able to model bifurcations in a state variable [9,29,45]. This model is starting to be more used in ecological studies and allows to detect three alternate dynamics of a system: linear, non-linear but still continuous, non-linear and discontinuous (i.e. regime shift behavior) [29,46,47]. The model was formulated by Thom in the 1970s and was enriched with a stochastic process (the Wiener process) in order to make it usable in natural systems [48,49]. The model is based on a cubic differential equation:

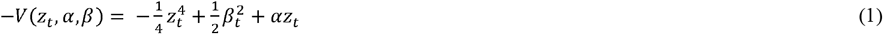

Where *V*(Z_*t*_, α, β) is a potential function, whose slope represents the rate of change of the system (the system state variable is called Z_*t*_), depending on the two control variables (α, β).

This equation is then extended with the Wiener process in order to render the model able to account for stochasticity:

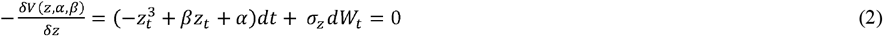

This equation models how a state variable (z_*t*_) abruptly changes depending on potential small changes of two control variables (α, β) [29,45,48,49]. In our case the state variable is modelled as a function of the cod population (CPUE), since our aim was to understand if all the three separate populations of the North Sea present discontinuous, regime shift dynamics (Eq.3a). Alpha is the so-called asymmetry parameter and controls the dimension of the state variable (if high or low). In our study alpha is a function of the fishing effort since fishing is believed to be the main controller of the dimension of the cod populations (Eq.3b). Beta is the so-called bifurcation parameter, that by changing can modify the relationship between the state variable and the asymmetry parameter from linear to non-linear discontinuous, and thus can induce hysteresis in the interaction between the state variable and the asymmetry variable. In our model beta was modelled as a function of the environmental drivers, the fish community and the big copepods resulting in many different models to compare (Eq.3c). The stochastic cusp model equation is then solved using the Cobb’s maximum likelihood method [48].

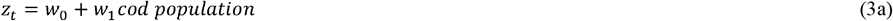

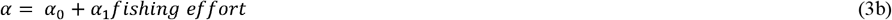

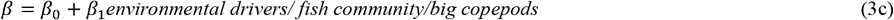

To detect the presence of discontinuous dynamics a Cardan’s discriminant (δ) needs to be solved (Eq.4):

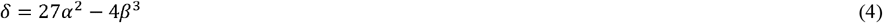

If the Cardan discriminant is smaller or equal to 0, then the state variable follows discontinuous dynamics and present multiple equilibria. Instead, if it is bigger, it presents linear dynamics. Thus, the model is able to detect different dynamics of the system [29,48,49]. The model is represented in 2D with a cusp plain, which is an area defined by the Cardan’s discriminant for which the combination of the drivers leads to a regime shift and thus to an area of very low resilience. The points of the state variable will move in and out of this area depending on which dynamics they are following. States of the system that are inside the cusp area (also called instability area or area below the fold) present instability and very low resilience.

The model automatically fits two alternative models, a linear model (Eq.5) and a logistic model (Eq.6) in order to directly do a model comparison and test which type of dynamic better suits the system.

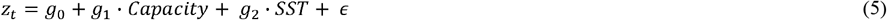

Where g_0_ is the intercept of the model, and g_1_ and g_2_ the slopes coefficients of the two control variables and ϵ is the normally distributed random error (mean=0, variance= σ square).

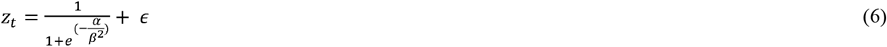

Where Z, α and β are canonical variables of the observed state and control variables defined in Eq.3 and ϵ is the zero mean random disturbance.

In order to compare the models, we used mainly the pseudo R squared calculated with the Cobb’s method and the AICc (Akaike Information Criterion corrected) just if needed since it is the selection based on the latter is still uncertain in this modelling approach. The selection of the models was performed to understand the prevalent dynamic of the three different populations of cod.

All the analyses were performed using R Studio Version 1.3.1093 (R Core Team, 2018). The change point analysis was conducted using the bcp and the changepoint packages [41,50]. The PCA was carried out using the ade4 and dudi.pca packages [51]. The cusp model was performed with the cusp package [49].

## Results

Using the NS-IBTS data we reconstructed time series of the CPUE of Atlantic cod in the 3 areas of the North Sea (South, North-West and Viking) where it is believed three different populations of cod reside (Fig.1 a). The time series of the CPUE of cod showed different dynamics in the three areas. Cod in the North-West North Sea showed a strong decline in CPUE around 1998 followed by relatively low abundances. After 2010, a recovery to almost pre-collapse levels occurred at the end of the time series (Fig.1b). A similar but more oscillatory pattern can be seen for cod in the Viking (North-East North Sea): cod declined around 1998 as well and remained low, with a small recovery towards the end of the time series (Fig.1c). In contrast, the dynamics of cod in the Southern North Sea appeared completely differently, with the CPUE values constantly declining without showing any sign of recovery (Fig.1d).

**Figure 1:**
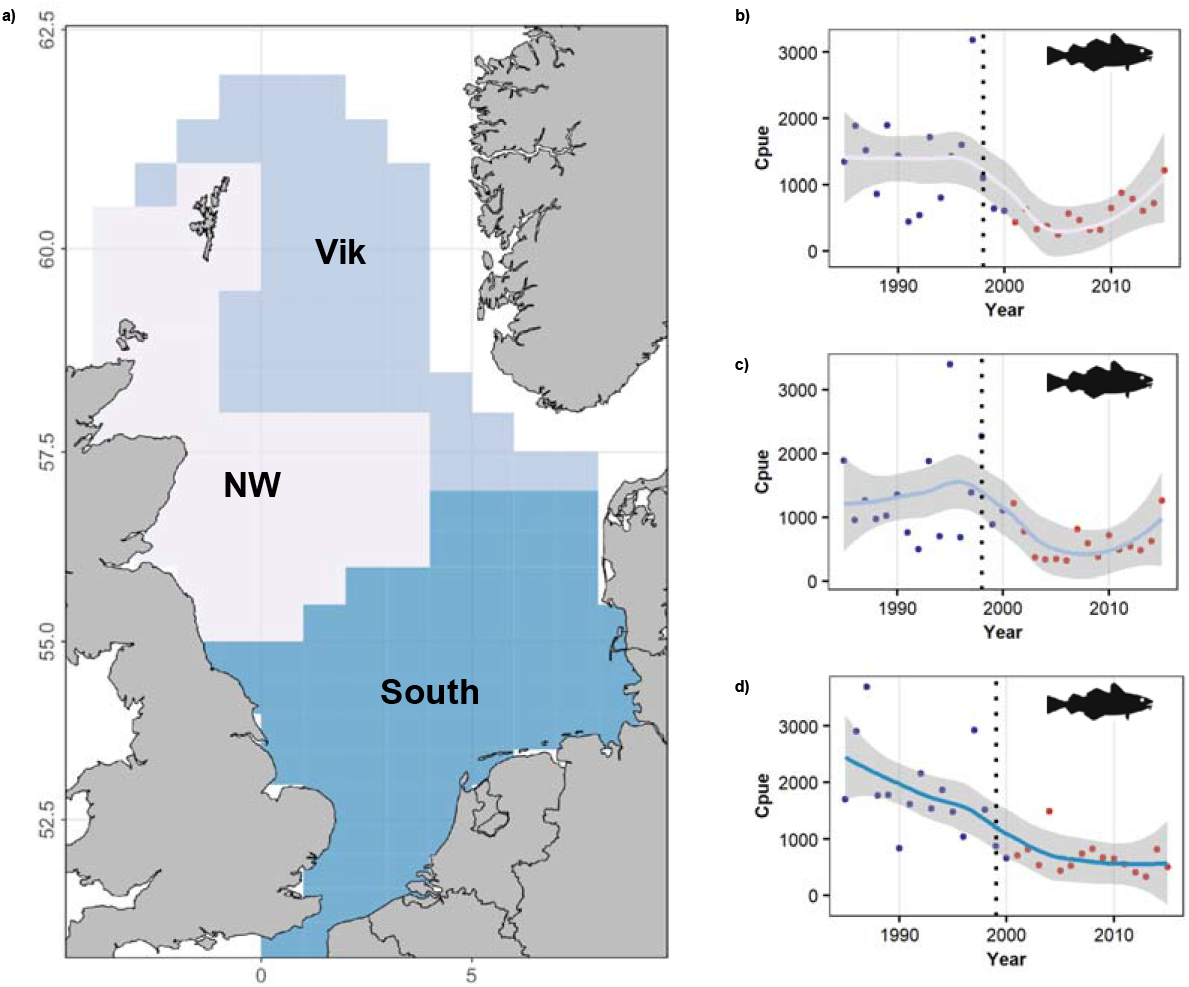
Time series of the three populations of cod in the North Sea. a) Map of the distribution of the three different cod populations in the North Sea as identified by the ICES Stock Assessment Group. Time series of CPUE of cod in b) The North-West North Sea, c) the Vikings, d) Southern North Sea. The different colors of the point correspond to the data before 2000 (blue) and after 2000 (red). The line passing through the dots is the smoothed trend done by fitting a loess to better visualize the trend of the populations. In grey is the confidence interval of the line. The vertical dotted line corresponds to the change point identified by the change point analysis (1998 for North-West and Viking cod and 1999 for Southern cod).

In order to identify abrupt shifts in the CPUE of cod that could indicate the presence of discontinuous dynamics, we performed a change point analysis. The three time series showed change points at a similar timing, 1998 or 1999 (dotted line Fig.1b-d) that led the CPUE of cod from a high to a low state. These results indicate that around the year 2000 cod populations abruptly changed in the entire North Sea, suggesting that regime shifts might have occurred.

To compare the dynamics observed in the cod populations with anthropogenic, biotic, and abiotic drivers we explored whether differences could be noted between the time series of the drivers in every area considered. The patterns were generally quite similar between North-West and Viking and in some cases a bit different for the Southern North Sea (Fig.2). For instance, while effort drastically declined in North-West and Viking over the time series (Fig.2a,b), it increased strongly in the South until 2000, and then started to decline (Fig.2c). Moreover, the magnitude of the effort, thus the number of hours of trawling was twice as high in the Southern North Sea compared to the Northern portion. In all the three areas copepods oscillated substantially, with the highest abundances present in the Viking and the lowest in the Southern North Sea (Fig.2d,e,f). Finally, temperature (both surface and bottom) and salinity showed similar dynamics in all the three areas (Fig.2g:o). However, the magnitude of temperature was higher in the Southern area.

**Figure 2:**
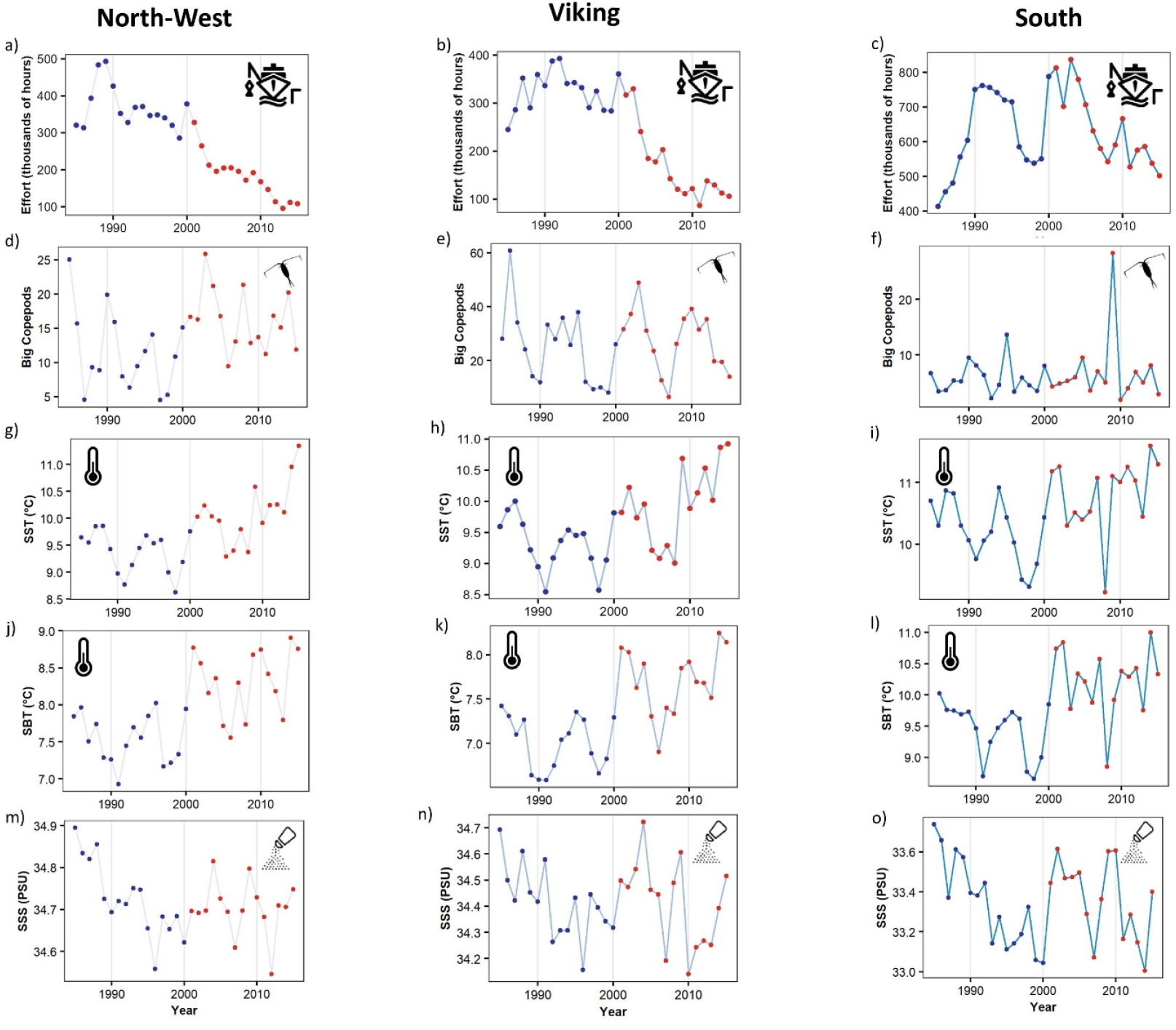
Anthropogenic biotic and abiotic drivers of cod in the three areas of the North Sea. Time series of effort (a, b, c), big copepods (d, e, f), Sea Surface Temperature (g, h, i) Sea Bottom Temperature (j, k, l) and Sea Surface Salinity (m, n, o) in the three areas (North-West, Vikings and South) of the North Sea. The color of the lines represents the areas as reported in Fig.1. The color of the dots represents the period before (blue) and after (red) 2000, the year in which the change points were detected.

To compare collapse and recovery patterns of cod populations to the entire fish community of the different areas, we performed a Principal Component Analysis on the fish communities of the three areas (SI A). The communities changed a lot over time, presenting a change point around the year 2000 (Fig.3). While the trends of the communities in all the three areas were similar, different species became dominant towards the end of the time series (Fig.3). In the North-West, cod and rays were abundant before 2000, while recently, sharks, gurnard, hake and flatfish became more common (Fig.3a, b). In the Viking area, the community trajectories were similar, except that rays increased towards the end of the time series (Fig.3c, d). Finally, in the Southern North Sea, cod and flatfish were dominant until 2000, while after 2000 gurnard and flatfish have become the most dominant species (Fig.3e, f). Thus, whereas some similarities can be found, differences are also clearly present in the community trajectories.

**Figure 2:**
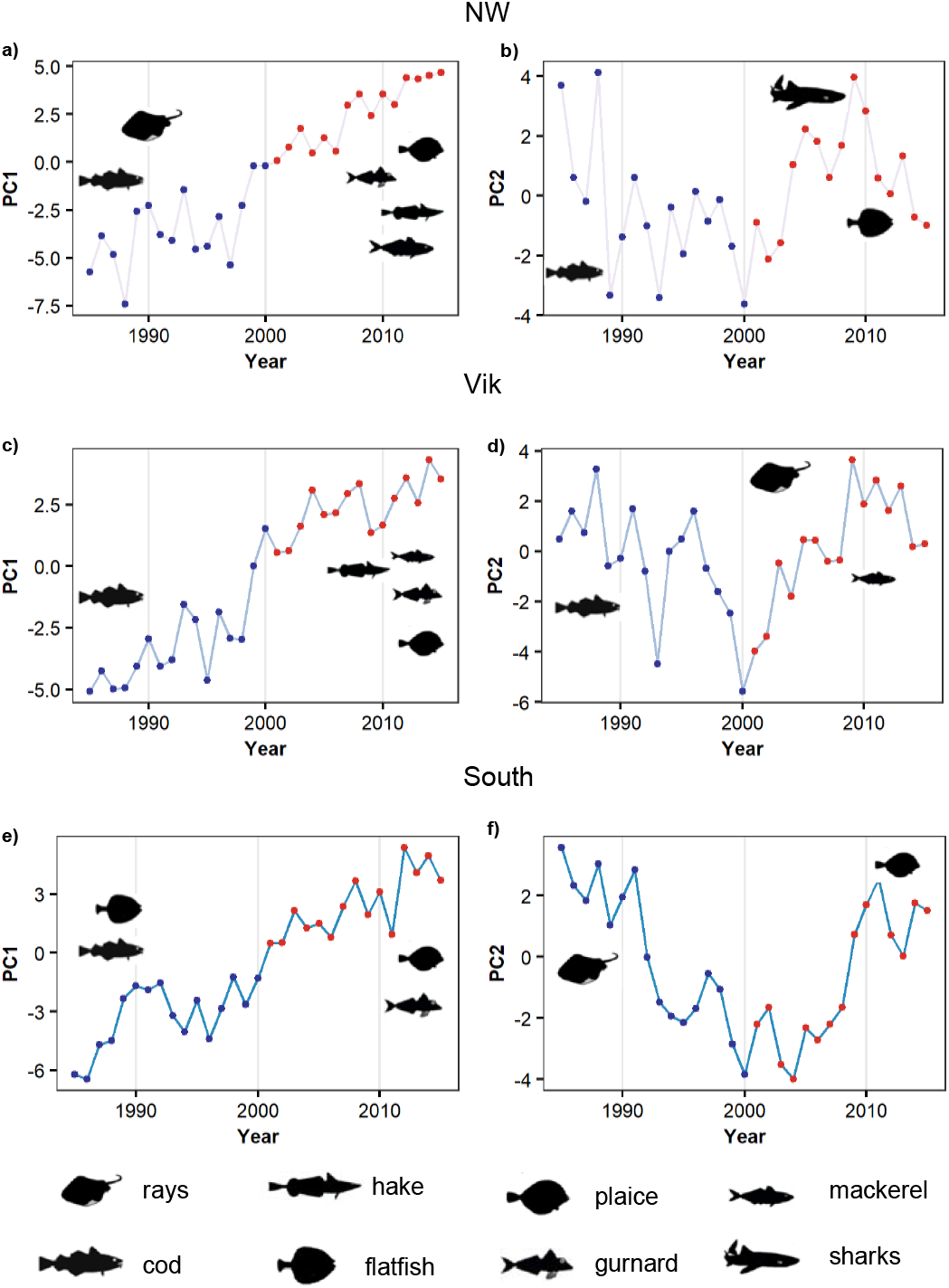
Community changes in the three areas of the North Sea. Changes in the community as highlighted by the Principal Component Analysis. PC1 and PC2 are used as proxy of the community status. PC1 and PC2 trends of the North-West North Sea community (a, b), the Viking community (c, d) and the Southern North Sea community (e, f). The color of the lines represents the areas as reported in Fig.1. The color of the dots represents the periods before (blue) and after (red) the change point in 2000. The icons show the dominant species for each of the two periods in every area.

After analyzing the possible drivers and determining the presence of abrupt shifts in the population time series, we analyzed hysteresis, a characteristic that could give a further indication of the possible presence of regime shift dynamics in the cod populations. The driver-state plots are the first step to do that and can indicate the presence of hysteresis in a state variable if, for the same level of the driver, multiple states of the state variable exist. We performed driver-state plots using fishing effort as the driver, since it is the driver directly influencing the CPUE of cod. The three areas showed different hysteresis patterns according to the state-driver plot (Fig.4). In the North-West a sign of hysteresis could be detected, but the loop was closed at the end of the time series when the level of CPUE reached past levels (Fig.4a). Instead, in the Viking and the Southern North Sea the hysteresis loop is not yet closed (CPUE levels lower compared to the beginning of the time series) and especially for the Southern North Sea the hysteresis is strong (Fig.4 b, c). These results could indicate that the cod populations in the three areas present a range of dynamics, with cod in the South and possibly Viking presenting regime shifts, while cod in the North-West presenting more linear and continuous dynamics. The information about the types of dynamics is fundamental to understand the different recovery patterns and potential among the three populations.

**Figure 4:**
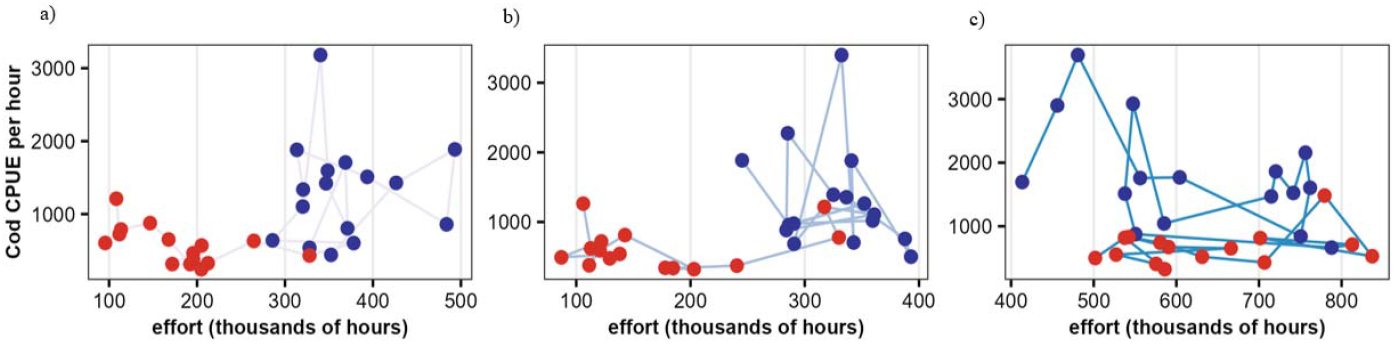
Hysteresis patterns of the three cod populations. The driver-state plot using as dependent variable CPUE of cod and as explanatory variable fishing effort for the a) North-West, b) Viking, c) Southern populations. The different colors of the point correspond to the data before 2000 (blue) and after 2000 (red).

To finally confirm the presence of different types of dynamics in the three cod populations of Atlantic cod in the North Sea, we performed a range of stochastic cusp models (SI B). Even though the model is a strong simplification of the reality, it can effectively help to confirm the presence of regime shifts in a state variable. For the three areas, we used as state variable the CPUE of cod, as asymmetry variable, the variable controlling the dimension of the state variable, fishing effort, and as bifurcation variable, the variable eventually inducing the regime shift, the various biotic and abiotic variables. The models returned pseudo R squared to evaluate the fit of the model for the three alternate models, the linear, the logistic and the cusp.

We thus selected the best R squared for each model. For the North-West North Sea the best model was always a logistic model, thus a continuous model (Tab.1, SI Tab.1, SI Fig.7). For the Viking, 4 out of 6 models were best fitted by a cusp, so there was almost a balance between continuous and discontinuous dynamics (Tab.1, SI Tab.2, SI Fig.8). Finally, 5 out of the 6 performed models for the Southern North Sea were best fitted through a cusp, thus with a model presenting a bifurcation (Tab.1, SI Tab.3, SI Fig.9). These results indicate that while cod in the North-West did not undergo regime shifts and present only continuous dynamics, cod in the Southern North Sea seemed to have undergone true discontinuous dynamics, and cod in the Viking seemed to have experienced a mixture of dynamics.

**Table 1:** Results of the stochastic cusp model. The table shows the different types of stochastic cusp models that were performed for the three areas. In particular the different bifurcation factors are indicated in the first column since the state variable and the asymmetry variable remained the same in all the models (CPUE and effort, respectively). For each area the best pseudo R squared among the three possible models, cusp, logistic and linear, are reported.

|  | NW | Vik | South |
| --- | --- | --- | --- |
| **SST** | Logistic 0.46 | Cusp 0.53 | Cusp 0.54 |
| **SSS** | Logistic 0.32 | Cusp 0.53 | Cusp 0.62 |
| **SBT** | Logistic 0.35 | Cusp 0.54 | Cusp 0.57 |
| **Big copepods** | Logistic 1 | Logistic 1 | Cusp 0.6 |
| **PC1** | Logistic 0.45 | Logistic 0.32 | Logistic 0.58 |
| **PC2** | Logistic 0.3 | Cusp 0.56 | Cusp 0.61 |

We then explored the results of the stochastic cusp model by plotting the cusp bifurcation space as modelled output (Fig.5). Since the output of the 10 different models were very similar for the same areas (See SI Fig.7,8,9), for simplicity, here we just show the results of the models using as bifurcation variable Sea Surface Temperature (SST). The output of the model shows if the system goes inside the instability area (blue area), the area in which three equilibria are possible. This area is the area of the transition thus the area that the system crosses during a regime shift. Systems that are inside this area present a very low resilience and are highly unstable [29,48]. The North-West North Sea cod was close to a tipping point (partially inside the bifurcation area) around the 1990s, but the immediate decline of fishing effort coupled with an increase in SST, pushed the stock away from the tipping point into a stable and resilient (far away from the bifurcation set) low to medium state (Fig.5 a). Therefore, this population did not undergo a regime shift. Cod in the Viking crossed a tipping point in 2000 (more points in the cusp area compared to cod in the North-West). Nonetheless, the population was able to stabilize, even if in a low state due to a decline of fishing effort and an increase of SST (Fig.5 b, red dots outside the cusp area). The difference between cod in the Viking and cod in the North-West North Sea is that in the former, discontinuous dynamics were highlighted (since generally the best model was a model with bifurcation), and therefore the population will likely never be able to return to the exact same state as before. Instead, since the North-West cod shows continuous dynamics, management actions could still manage to bring the population back towards past abundances. Finally, cod in the Southern North Sea showed a very different situation. Here the system presented always a low resilience going in and out from the transition area (the blue area) and is at the moment really unstable and locked in a low state (Fig.5 c, small red dots). Thus, for this stock no return to previous condition seems possible since the stock underwent a regime shift and its resilience is very low.

**Figure 5:**
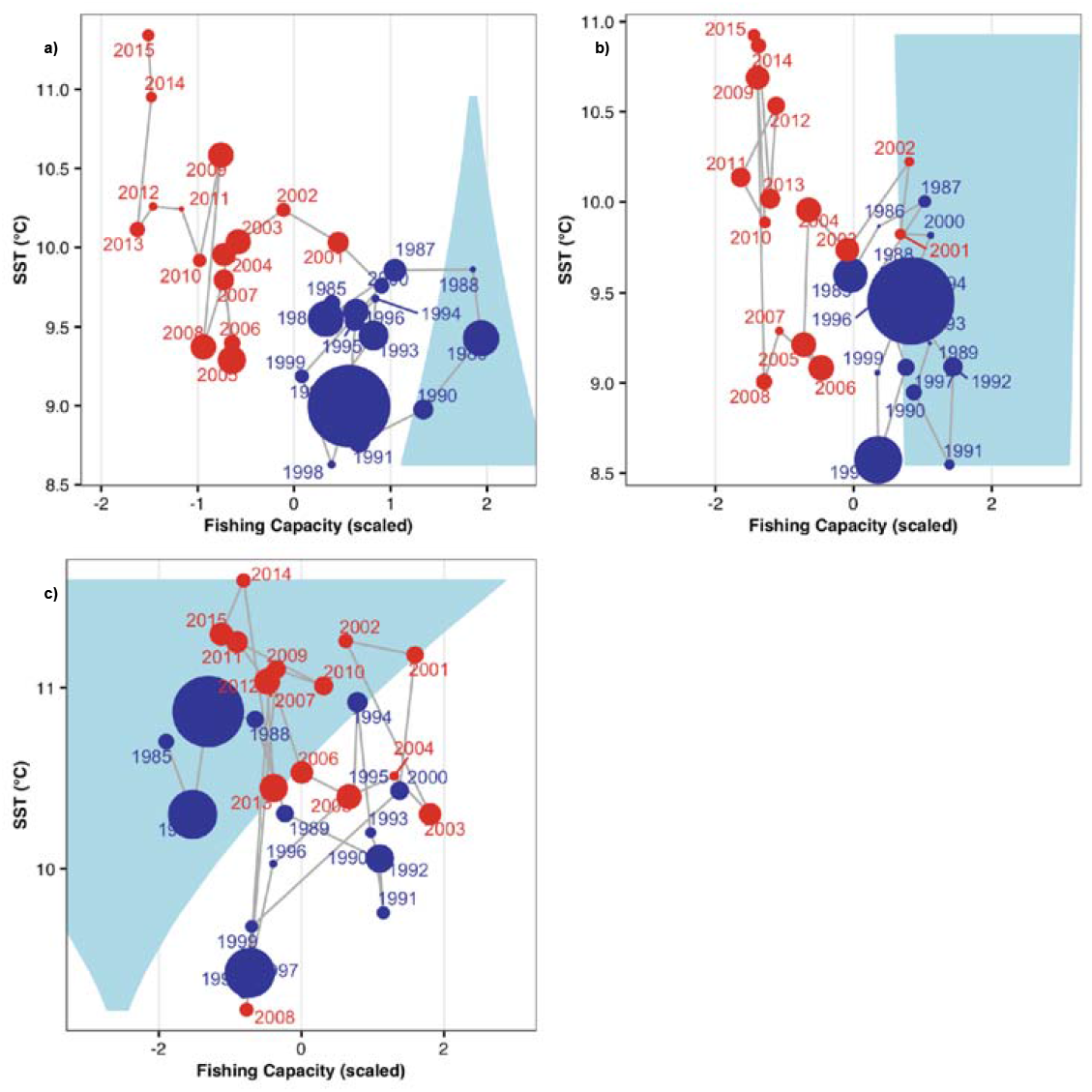
The stochastic cusp models. Output of the stochastic cusp models for a) the North-West North Sea cod, b) the Viking cod and c) the Southern cod. On the x axis the asymmetry variable, fishing capacity expressed in hours and scaled. On the y axis the bifurcation factors in this case Sea Surface Temperature (SST). The blue area corresponds to the transition area, the area where three equilibria of the systems are present: two stable and one unstable. The dimension of the dots is proportionate to the CPUE of the cod population (big dot = high CPUE, small dots = low CPUE). The colors of the points represent the periods (blue) before and (red) after 2000.

## Discussion

In this study we applied a range of techniques, from change point models and state-driver plots to the stochastic cusp model to investigate whether regime shifts can be found in three putative populations of Atlantic cod in the North Sea depending on a variety of external drivers. This study explores an urgent question, because the stock of Atlantic cod in the North Sea has been considered as a single unit from a management point of view until very recently [14,32–34]. In the last years, the management unit has undergone a review process and it was proposed to split it into three units, consistent with genetics and ecological evidence, in the attempt to provide more accurate management measures [30,36]. A clear understanding of the presence of regime shifts in the three units however, could not yet be determined in a structural way [36]. Our results clearly and unequivocally show that the three populations of cod indeed experience contrasting dynamics. While the North-West population in our analyses presents non-linear but continuous dynamics (i.e. not regime shifts), the Viking and the Southern cod show regime shift dynamics. This is fundamental from a management point of view since it indicates that different recovery patterns might be found among the three putative populations. If effective management is applied, the North-West population could be able to return to previous abundance: the continuous dynamics indicate that if the driver of change is reverted, the population can potentially return to the past state [10,17]. At the moment, in fact, the North-West population shows encouraging signs of recovery [30,31,52,53].

Very different is instead the fate of the Viking and Southern populations. These two units present discontinuous dynamics and regime shifts, thus hysteresis hinders the recovery towards the previous state, and the populations will likely be trapped in a depleted state in the future. Still, important differences can be found between the two populations. The Viking cod is in a high resilience state: this means that its dynamics are expected to be more stable, and thus easier to manage in order to maintain it at low but stable levels. Therefore, if exploited at sustainable levels, the Viking population should be able to remain robust and stable in the face of other drivers. Instead, cod in the Southern North Sea shows a very low resilience to external drivers and thus is more unstable and more difficult to manage. The population in the Southern North Sea is in a very depleted state and management measures may result ineffective [31,53,54]. Notably, these results complement and advance the analysis on regime shifts in the recruitment productivity of the three sub-populations performed at the latest Benchmark meeting [36] to identify the emergence of novel recruitment regimes. A regime shift in 1997 was shown using change point analyses for the three sub-populations, much stronger for the Southern North Sea cod. Our detailed investigation on the drivers and the underlying mechanisms of the regime shifts following the methodological guidelines of Sguotti et al (2022), confers a more robust understanding of the occurrence of regime shifts in the three populations, showing that the North-west population did not undergo a regime shift.

Even if the three populations show contrasting patterns, the trends of the drivers used as bifurcation factors and impacting them are rather similar. Indeed, similarities can be seen in the trends of temperature (both surface and bottom) with steady increase, but also in the time series of copepods and salinity. The magnitude of the drivers is however different. The average sea surface temperature in the Southern North Sea is 2°C higher compared to temperature in the Northern areas. This indicates that cod in the Southern North Sea is exposed to higher temperatures that are closer to the maximum thermal tolerance of the species [54–58]. Temperature can directly act on cod through changes in recruitment or growth, or indirectly by changing the time or occurrence of their preys [29,55,58–60]. Therefore, the environment in the Southern North Sea is more hostile for cod compared to the other areas [31,61]. Instead, temperature in the Northern North Sea is acceptable for cod, allowing it to thrive and recover [57,58]. The quantities of big copepods are also lower in the Southern North Sea compared to the other areas. Being big copepods fundamental prey for cod larvae, the low abundances might represent another limiting factor in the South [26].

Apart from changes in biotic and abiotic drivers, also changes in the community display major differences among the three areas. While the communities in the Viking and North-West North Sea are still dominated by some gadoids, the Southern North Sea is now dominated by plaice, gurnard, and forage fish. Many of these species are strong predators of young cod and recruits at the settlement stage, and therefore their increase can render the Southern North Sea environment more hostile for cod settlers and juveniles, hindering the recovery towards previous abundances [59,61–64]. As a result, the population of cod might soon disappear from this area. Fishing pressure is the other driver that clearly presents different trends among the three areas. Indeed, fishing intensity has been higher in the Southern North Sea for centuries and this could have chronically impacted fish populations since longer times, again causing the southern population to be less resilient from other stressors [40,65]. Similar patterns of recovery and disappearance can be found in shark and ray populations, that are absent or extirpated from the Southern North Sea but are showing recovery in the Northern areas [52,66,67], arguably because of the difference in fishing pressure in the two areas. Thus, past fishing trends could have a strong effect on fish populations dynamics even in the present, and, coupled with environmental drivers and community changes, cause the presence of discontinuous dynamics at lower latitudes in the North Sea.

Our study presents some limitations that need to be taken into account when interpreting the results. First of all, the resolution of the putative populations of cod in the North Sea used in our study is not univocal. The latest cod stock benchmark [36] recognizes connectivity between the North-West area and the West of Scotland population, which we have not included in our study. Our distribution follows the North Sea stock subdivision that has been proposed in earlier assessments, limited to North Sea proper. While it could be speculated that this mismatch may explain the difference observed (e.g. the observed significant-but small-regime shift in ICES Benchmark for this population), we propose that such differences may rather be driven by the more detailed analysis performed in our study. This effectively highlights the importance of exploring results with diverse methods. Regardless, the main conclusions of our study, showcasing the emergence of spatial regime shifts, is not impacted by the inclusion or not of the West of Scotland stock. A further confirmation of the presence of spatial regime shift and the importance of scale derives from consideration about alternative population resolution: in fact, it has been debated whether the Southern and North-West populations should be consider separately or as one unit, given their genetic homogeneity [30,34]. For this reason, we repeated the analyses also considering just one population instead of two (i.e. merging these two units), and we showed that the emergent dynamics are masked if we aggregate these populations together since the two areas present opposite dynamics (SI C). Indeed, the large population derived from the aggregation of the two sub-populations presents no recovery, strong hysteresis, and discontinuous dynamics (SI C). This indicates that the dynamics of the large population match the dynamics of the Southern cod, but completely mask the good status of the North-West cod. The use of three populations, matching the recognized structure for assessment, allows to better highlight and unmask contrasting dynamics in the populations that might be useful for management.

Another bias lies in the fact that fishing pressure is the results of hours of fishing of beam and otter trawls [40]. Cod is not targeted by beam trawls and is also not always the target of otter trawls. However, using this non-specific fishing pressure, we capture both, direct (extraction) and indirect effects of fishing such as bycatch and habitat destruction which can impact the cod populations [68,69]. Finally, some inherent bias resides in the use of the stochastic cusp model. Indeed, this model, as generally statistical models, entails a simplification of reality, but can nonetheless provide important information about contrasting dynamics [9,29]. Moreover, we applied a number of techniques to detect regime shifts (from change point to state-driver plots) and the cusp model confirmed and better explained the results obtained with the other methods. It is still unclear whether the AICc could be used to perform model comparison, therefore we used the R squared [29,49]. Thus, we believe that considering these possible biases of the analyses, the results clearly support our interpretation.

How can these results be useful for management? Here we show, aligning with the existing literature but from a novel point of view and approach, that considering just one stock of Atlantic cod for the whole North Sea might lead to incorrect assumptions which may result in inappropriate management measures since the three sub-populations of cod show contrasting dynamics and recovery potential. Our results align with, and support the conclusions of, the latest ICES benchmark (2023). We highlight that separated management should be applied in the three areas considering that, based on our analysis, the North-West cod could potentially recover to previous abundances, Viking cod could be managed but cannot go back to previous levels and Southern cod appears to be really difficult to manage and recover. These dynamics could also create conflict among the fishers, potentially inducing spatial redistribution of fisheries from the Southern part of the North Sea to the northern, or changes of target species [70–72].

More in general, our study highlights the importance of looking at regime shifts from a spatial point of view, and how heterogeneous dynamics can emerge at different spatial scales [12,14]. It shows the difficulty of detecting regime shifts especially in large marine ecosystems that can present a variety of habitats, communities, and anthropogenic impacts. More research is needed to understand how to tackle the problem of spatial scales in regime shift science and to understand which scales are meaningful to use [15]. Our analyses revealed that ecologically units are not necessarily meaningful management units and that statistical analyses can help to unravel important patterns that might be useful for management. Understanding regime shifts is important to apply meaningful and effective management measures [16–18,73]. New avenues should be pursued to make regime shifts science more useful for management and to integrate it in policy frameworks [15].

## Acknowledgments

This study was conducted within the course of the BMBF-funded project SeaUseTip, Spatio-temporal analysis of tipping points in the socio-ecological system of the North Sea (funding code: 01LC1825A-C), and benefitted from exchanges and discussion at the ICES WGCOMEDA. EC was funded with the support of the Erasmus+ programme of the European Union. We thank the scientists that collected the data that were used for this study.

## Author’s contribution

CS, GR and CM had the idea and designed the study, EC and CS performed the analyses and produced the figures, all the authors helped with the writing of the manuscript.

## Data availability statement

The data are available online in the websites provided in Material and Methods.

