## Supplementary Information (SI) for "Evidence of spatial regime shifts: contrasting spatial dynamics of three Atlantic cod (*Gadus* morhua) populations in the North Sea"

**Supporting Information A: the PCA Analyses**

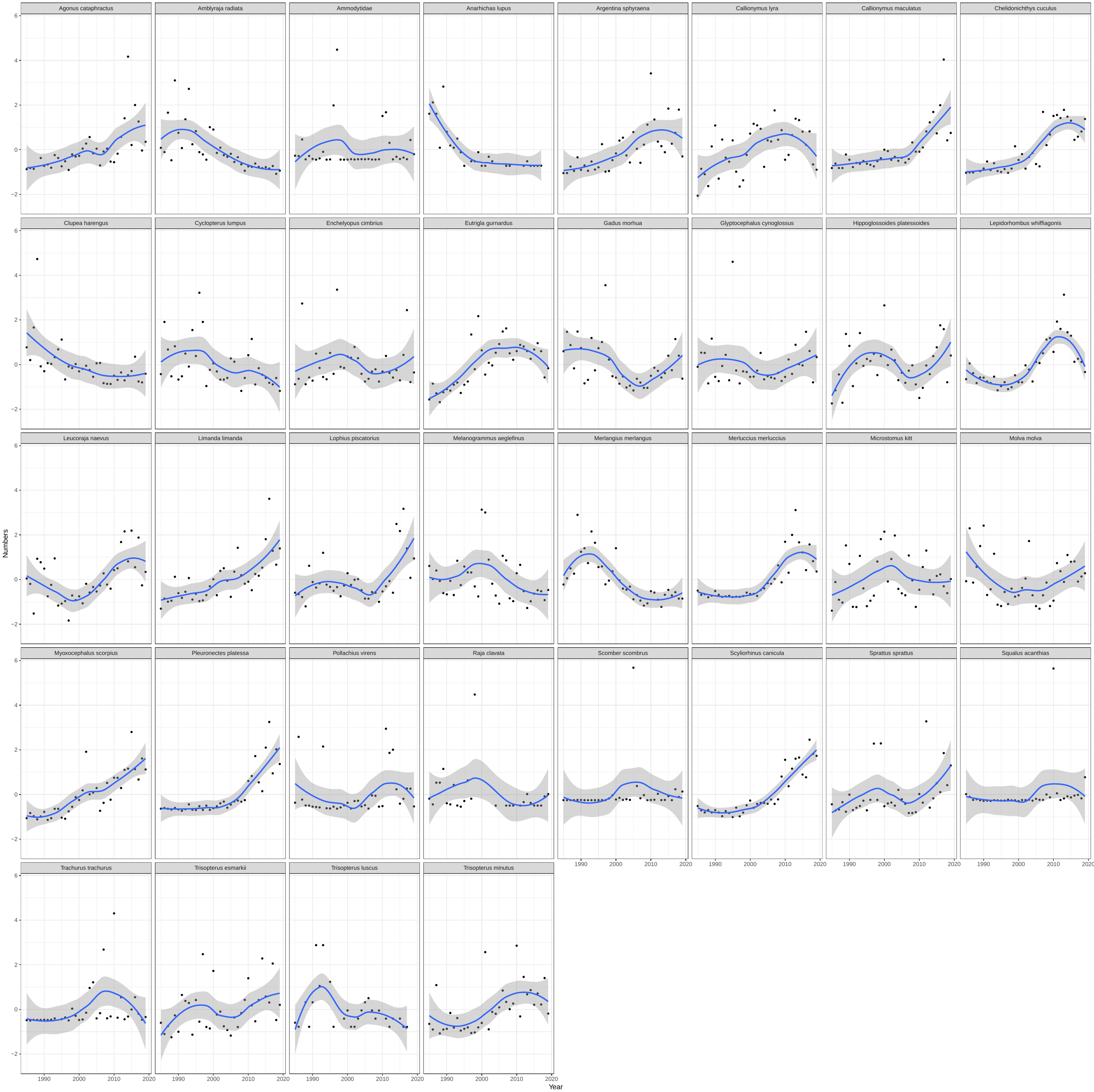

**Figure 1:** **Species used for the PCA of the North-West North Sea community**. Time series of the average yearly abundance of the species in the North-West North Sea. The time series were built from the NS-IBTS data and go from 2015 to 2020. The blue line is the smoothed trend (with the confidence interval in grey).

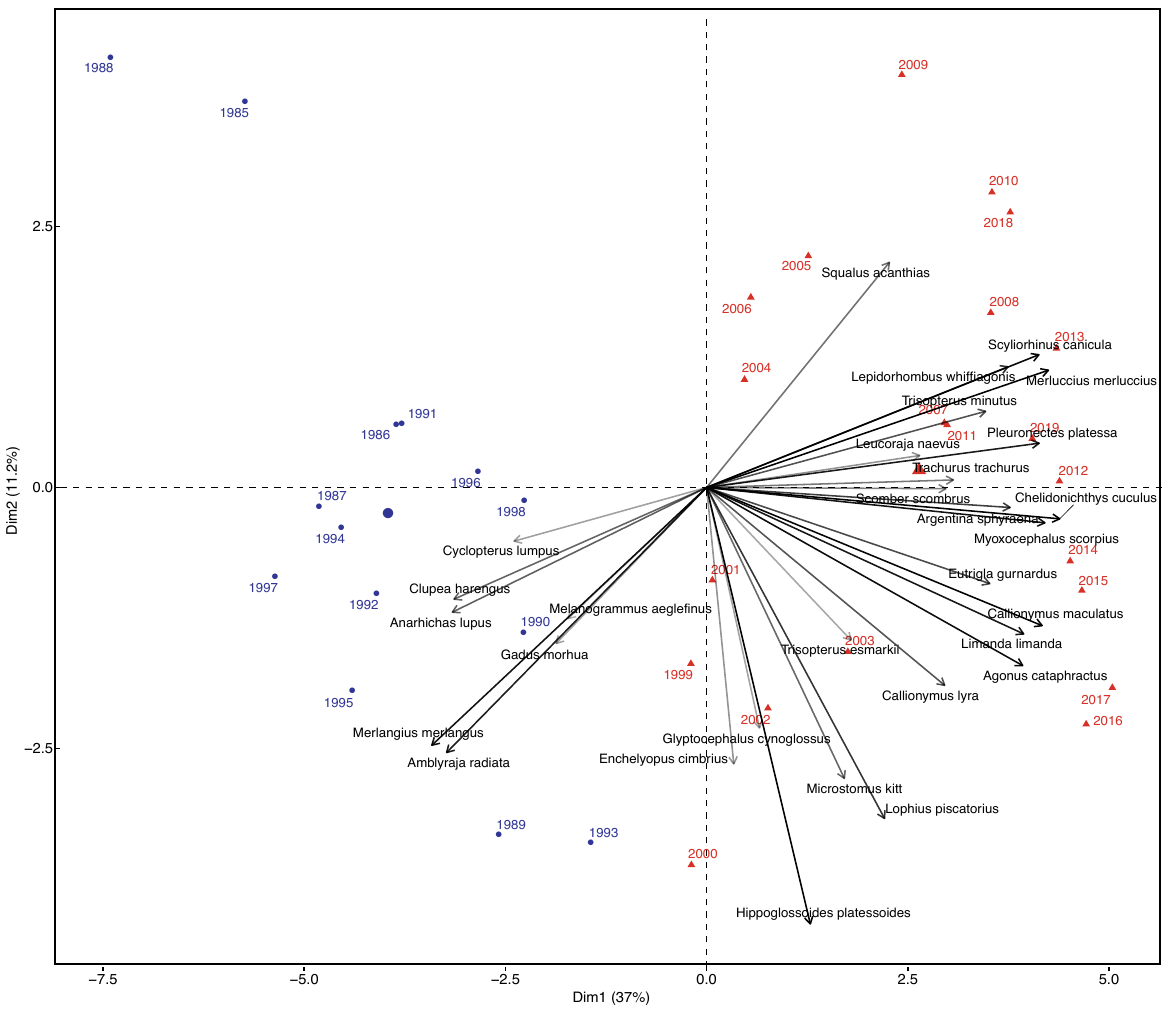

**Figure 2: PCA of the North-West North Sea community**. PC1 and PC2 explain respectively 37% and 11.2% of the variance. The years are colored by the clusters identified by the hierarchical clustering analysis (in blue before 2001).

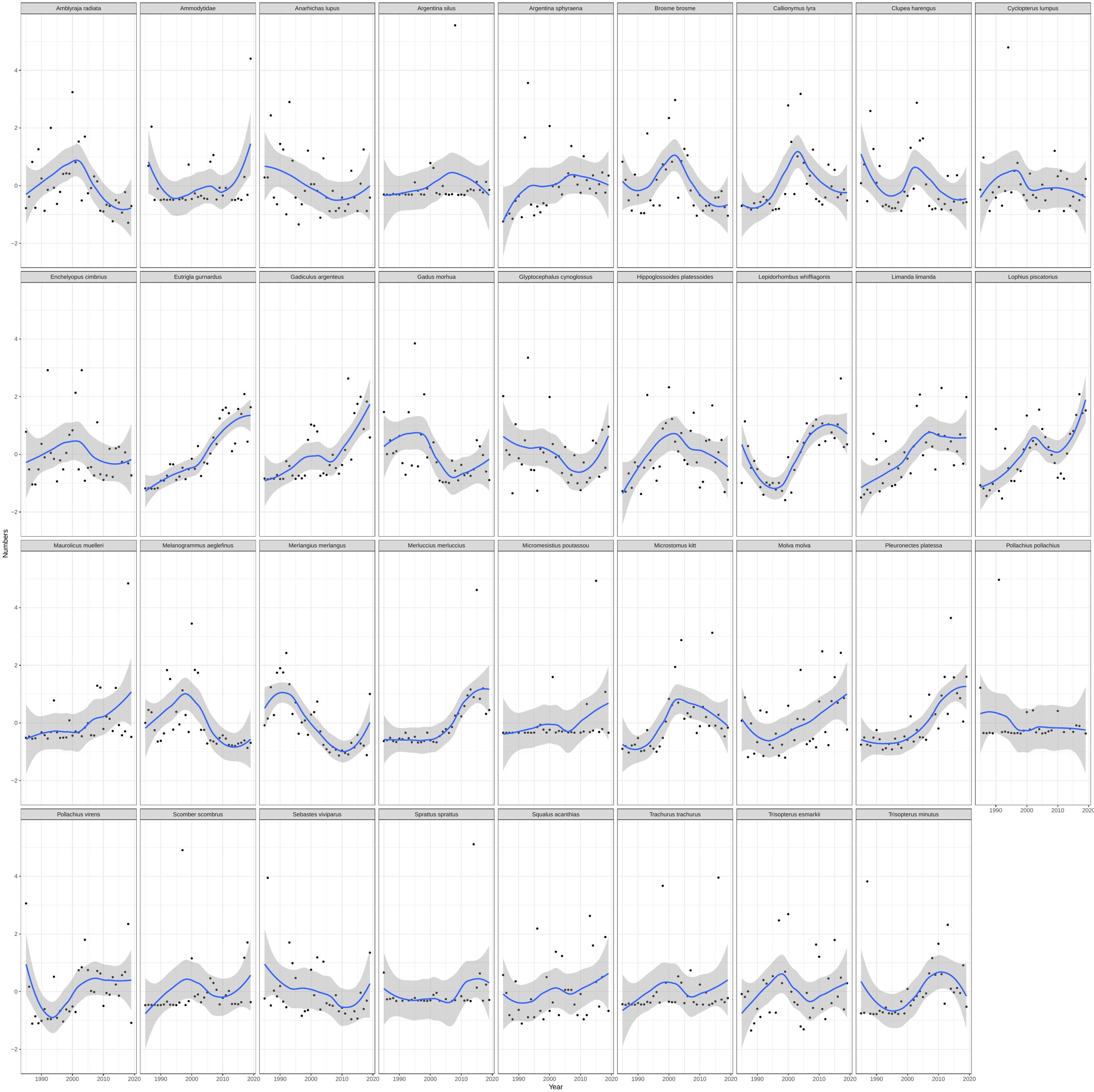

**Figure 3: Species used for the PCA of the Viking North Sea community**. Time series of the average yearly abundance of the species in the Viking North Sea. The time series were built from the NS-IBTS data and go from 2015 to 2020. The blue line is the smoothed trend (with the confidence interval in grey).

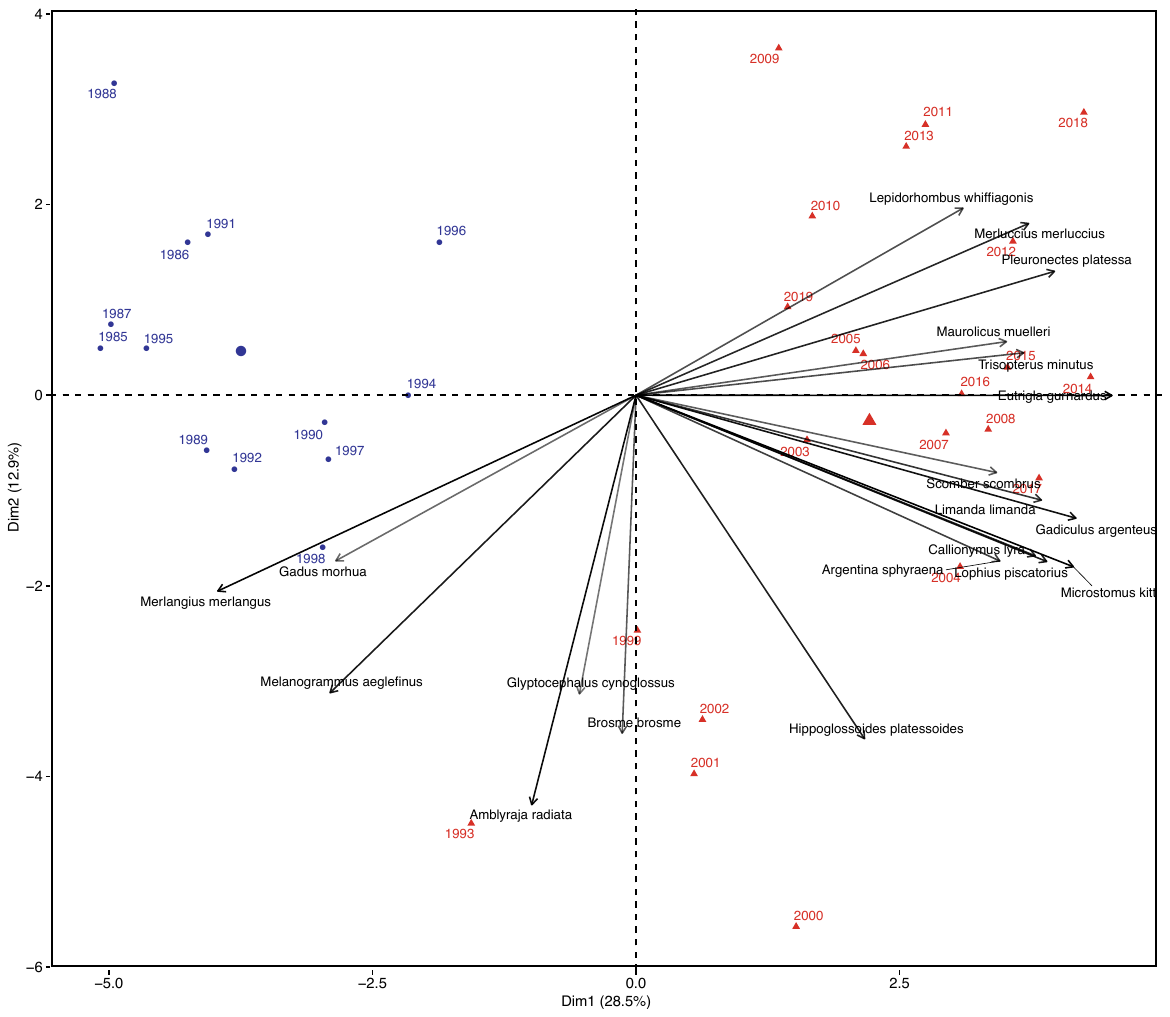

**Figure 4: PCA of the Viking North Sea community**. PC1 and PC2 explain respectively 28.5% and 12.9% of the variance. The years are colored by the clusters identified by the hierarchical clustering analysis (in blue before 2001).

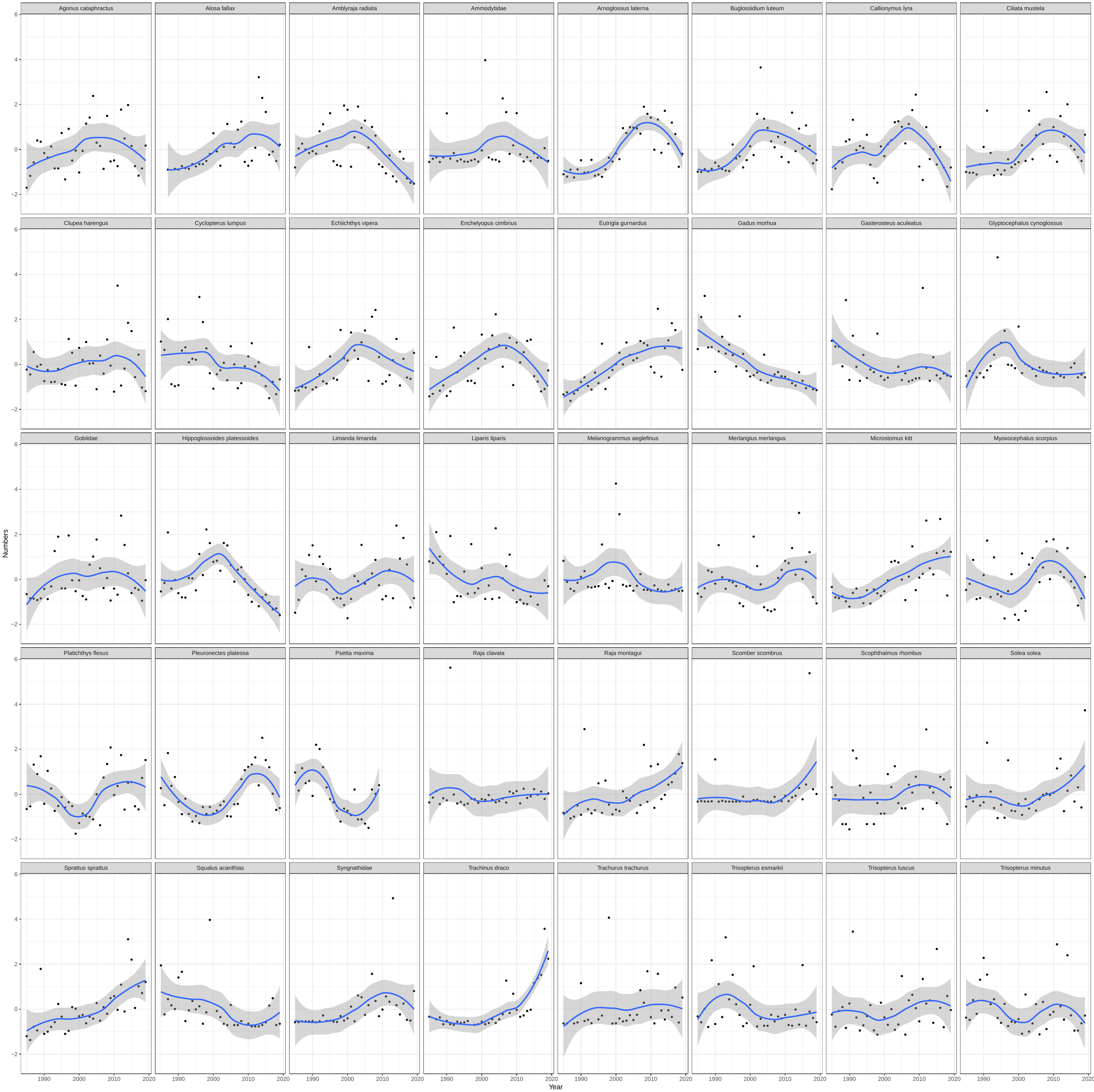

**Figure 5: Species used for the PCA of the Southern North Sea community**. Time series of the average yearly abundance of the species in the Southern North Sea. The time series were built from the NS-IBTS data and go from 2015 to 2020. The blue line is the smoothed trend (with the confidence interval in grey).

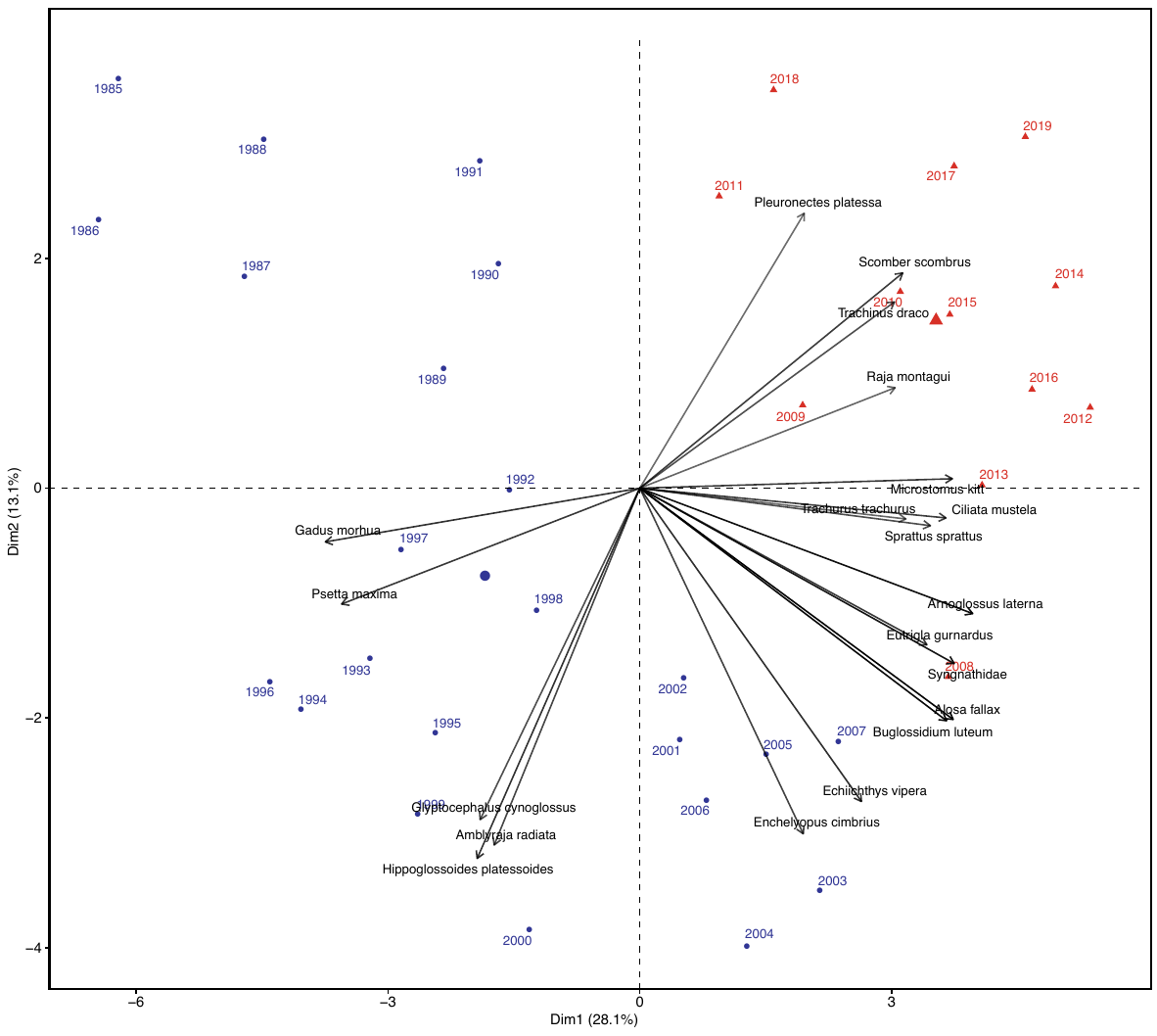

**Figure 6: PCA of the Southern North Sea community**. PC1 and PC2 explain respectively 28.1% and 13.1% of the variance. The years are colored by the clusters identified by the hierarchical clustering analysis (in blue before 2001).

**Supporting Information B: the stochastic cusp model**

**Table 1. Results of the cusp models for the North-West North Sea cod.** Estimates of the parameters of the stochastic cusp model. α1 and α2 are respectively the intercept and slope of the splitting parameter depending on fishing. β1 and β2 are respectively the intercept and slope of the bifurcation parameter *that changes in all the models*. w1 and w2 are the intercept and the slope of the state variable*, the abundance of cod*. The significant levels of the estimates are indicated with a star (* p < 0.05, ** p< 0.005, *** p< 0.0005). The R^2^ are pseudo Cobb R^2^ computed to compare the model using the cusp formulation or a linear *or a logistic* model*.*

| **Parameters** | **SST** | **SSS** | **SBT** | **Big copepods** | **PC1** | **PC2** |
| --- | --- | --- | --- | --- | --- | --- |
| **α1** | -1.85* | -1.78* | -1.80* | -1.83* | -1.47* | -1.76* |
| **α2** | 1* | 0.84* | 0.85* | 0.78 | 0.43 | 0.78 |
| **β1** | 6.57 | 29.9 | 1.13 | 0.37 | 2.04* | 1.38 |
| **β2** | -0.58 | -0.82 | 0.01 | 0.08 | 0.16 | 0.18 |
| **w1** | -1.14*** | -1.22*** | -1.22*** | -1.37*** | -1.4*** | -1.28*** |
| **w2** | 0.78*** | 0.76*** | 0.75*** | 0.72*** | 0.74*** | 0.76*** |
| **R^2^ linear** | 0.21 | 0.21 | 0.21 | 0.29 | 0.26 | 0.23 |
| **R^2^ logistic** | 0.45 | 0.32 | 0.35 | 1 | 0.45 | 0.33 |
| **R^2^ cusp** | 0.07 | 0.17 | 0.17 | 0.25 | 0.23 | 0.21 |

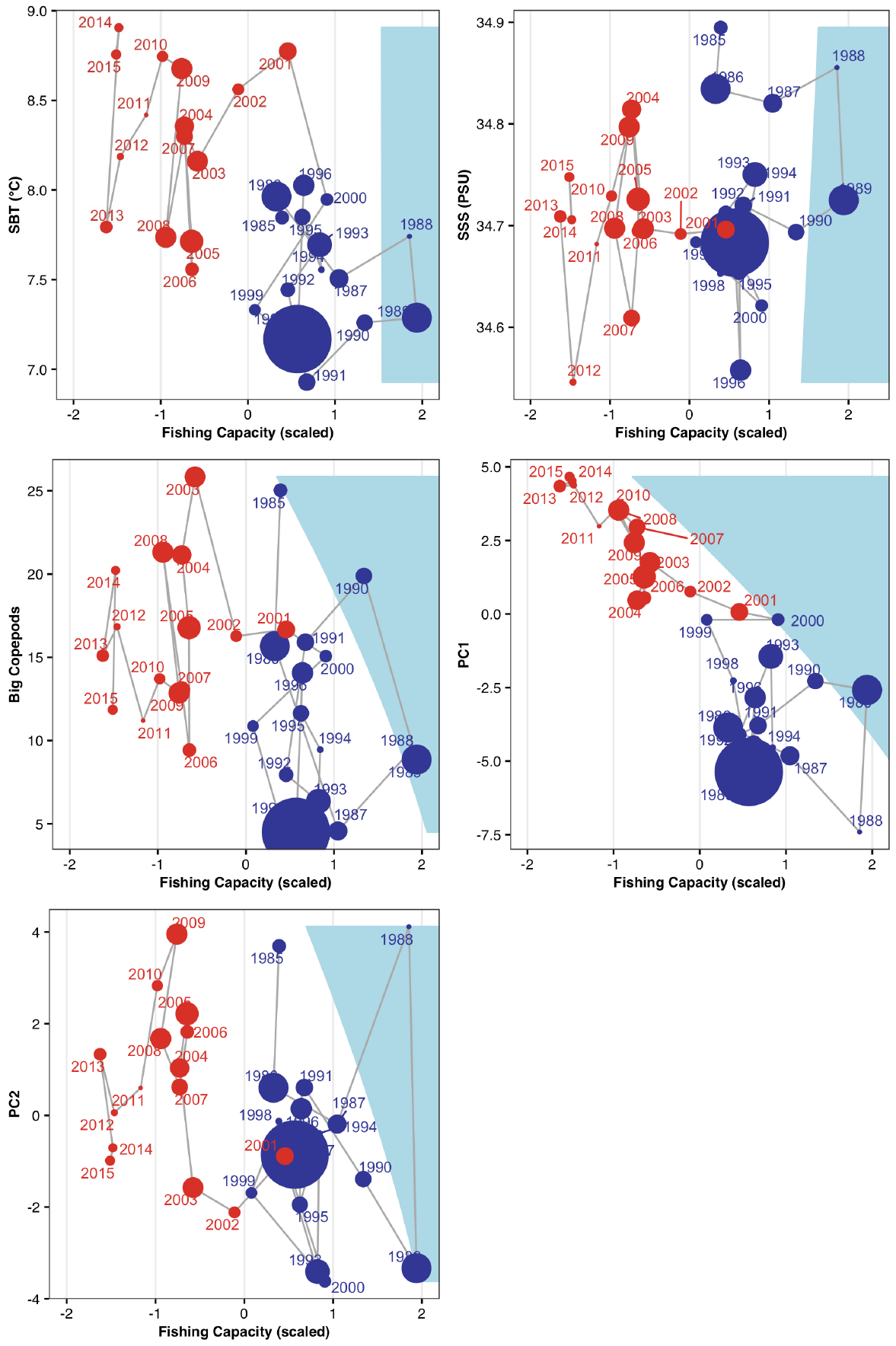

**Figure 7: The stochastic cusp models of the North-West North Sea cod.** Output of the stochastic cusp models with the different beta variables. On the x axis the asymmetry variable, fishing capacity expressed in hours and scaled. On the y axis the bifurcation factors as linear models of the different biotic and abiotic variables. The blue area corresponds to the transition area, the area where three equilibria of the systems are present; two stable and one unstable. The dimension of the dots is proportionate to the CPUE of the cod population (big dot = high CPUE, small dots = low CPUE). The colors of the points represent the periods (blue) before and (red) after 2000.

**Table 2. Results of the cusp models for the Viking North Sea cod.** Estimates of the parameters of the stochastic cusp model. α1 and α2 are respectively the intercept and slope of the splitting parameter depending on fishing. β1 and β2 are respectively the intercept and slope of the bifurcation parameter that changes in all the models. w1 and w2 are the intercept and the slope of the state variable, the abundance of cod. The significant levels of the estimates are indicated with a star (* p < 0.05, ** p< 0.005, *** p< 0.0005). The R^2^ are pseudo Cobb R^2^ computed to compare the model using the cusp formulation or a linear or a logistic model.

| **Parameters** | **SST** | **SSS** | **SBT** | **Big copepods** | **PC1** | **PC2** |
| --- | --- | --- | --- | --- | --- | --- |
| **α1** | -1.61* | -1.64** | -1.54* | -1.61** | -1.13* | 0.01* |
| **α2** | 0.83 | 0.85* | 0.76 | 0.87* | 0.21 | 0.63 |
| **β1** | 1.34 | 13.3 | 1.16 | 1.21 | 2.99*** | 2.22** |
| **β2** | 0.06 | -0.33 | 0.25 | 0.02 | 0.27* | 0.21 |
| **w1** | -1.38*** | -1.38*** | -1.38*** | -1.36*** | -1.58*** | -14*** |
| **w2** | 0.8*** | 0.8*** | 0.8*** | 0.82*** | 0.81*** | 0.83*** |
| **R^2^ linear** | 0.16 | 0.16 | 0.16 | 0.16 | 0.29 | 0.16 |
| **R^2^ logistic** | 0.29 | 1 | 1 | 0.29 | 0.32 | 0.30 |
| **R^2^ cusp** | 0.53 | 0.53 | 0.54 | 0.55 | 0.26 | 0.57 |

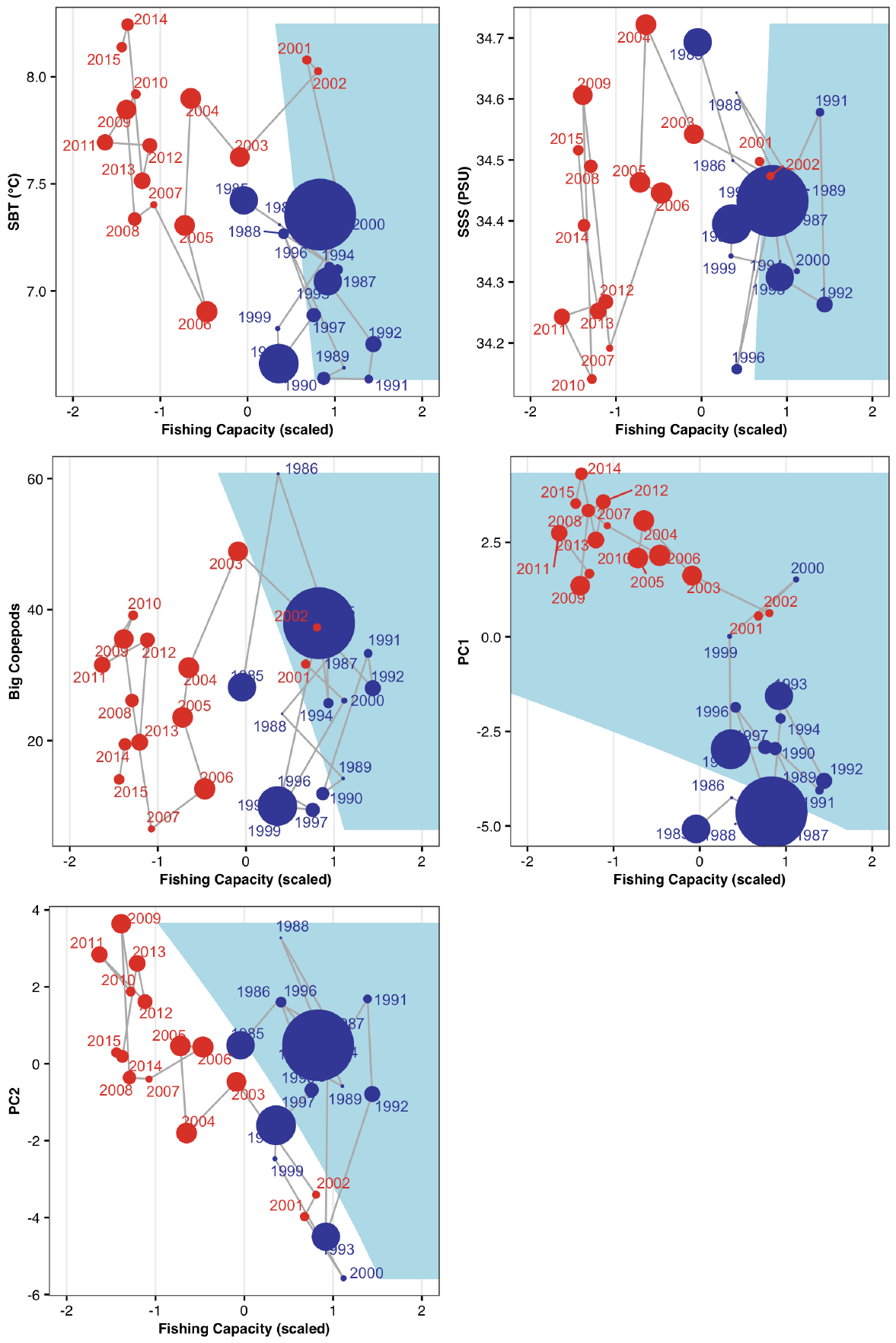

**Figure 8: The stochastic cusp models of the Viking cod.** Output of the stochastic cusp models with the different beta variables. On the x axis the asymmetry variable, fishing capacity expressed in hours and scaled. On the y axis the bifurcation factors as linear models of the different biotic and abiotic variables. The blue area corresponds to the transition area, the area where three equilibria of the systems are present; two stable and one unstable. The dimension of the dots is proportionate to the CPUE of the cod population (big dot = high CPUE, small dots = low CPUE). The colors of the points represent the periods (blue) before and (red) after 2000.

**Table 3. Results of the cusp models for the Southern North Sea cod.** Estimates of the parameters of the stochastic cusp model. α1 and α2 are respectively the intercept and slope of the splitting parameter depending on fishing. β1 and β2 are respectively the intercept and slope of the bifurcation parameter that changes in all the models. w1 and w2 are the intercept and the slope of the state variable, the abundance of cod. The significant levels of the estimates are indicated with a star (* p < 0.05, ** p< 0.005, *** p< 0.0005). The R^2^ are pseudo Cobb R^2^ computed to compare the model using the cusp formulation or a linear or a logistic model.

| **Parameters** | **SST** | **SSS** | **SBT** | **Big copepods** | **PC1** | **PC2** |
| --- | --- | --- | --- | --- | --- | --- |
| **α1** | -0.97** | -0.96* | -1.00* | -0.95** | -1.09* | -0.95* |
| **α2** | -0.37 | -0.34 | -0.35 | -0.33 | -0.37 | -0.31 |
| **β1** | -10.5 | 53.1 | -11.6* | 1.54* | 3.22*** | 1.64* |
| **β2** | 1.17* | -1.54 | 1.37** | 0.013 | 0.55*** | -0.04 |
| **w1** | -1.08*** | -1.04*** | -1.16*** | -1.02*** | -1.57*** | -1.03*** |
| **w2** | 0.94*** | 0.91*** | 0.93*** | 0.911*** | 0.94*** | 0.91*** |
| **R^2^ linear** | 0.15 | 0.1 | 0.22 | 0.08 | 0.55 | 0.09 |
| **R^2^ logistic** | 0.42 | 0.4 | 0.49 | 0.37 | 0.58 | 0.37 |
| **R^2^ cusp** | 0.54 | 0.62 | 0.58 | 0.61 | 0.51 | 0.61 |

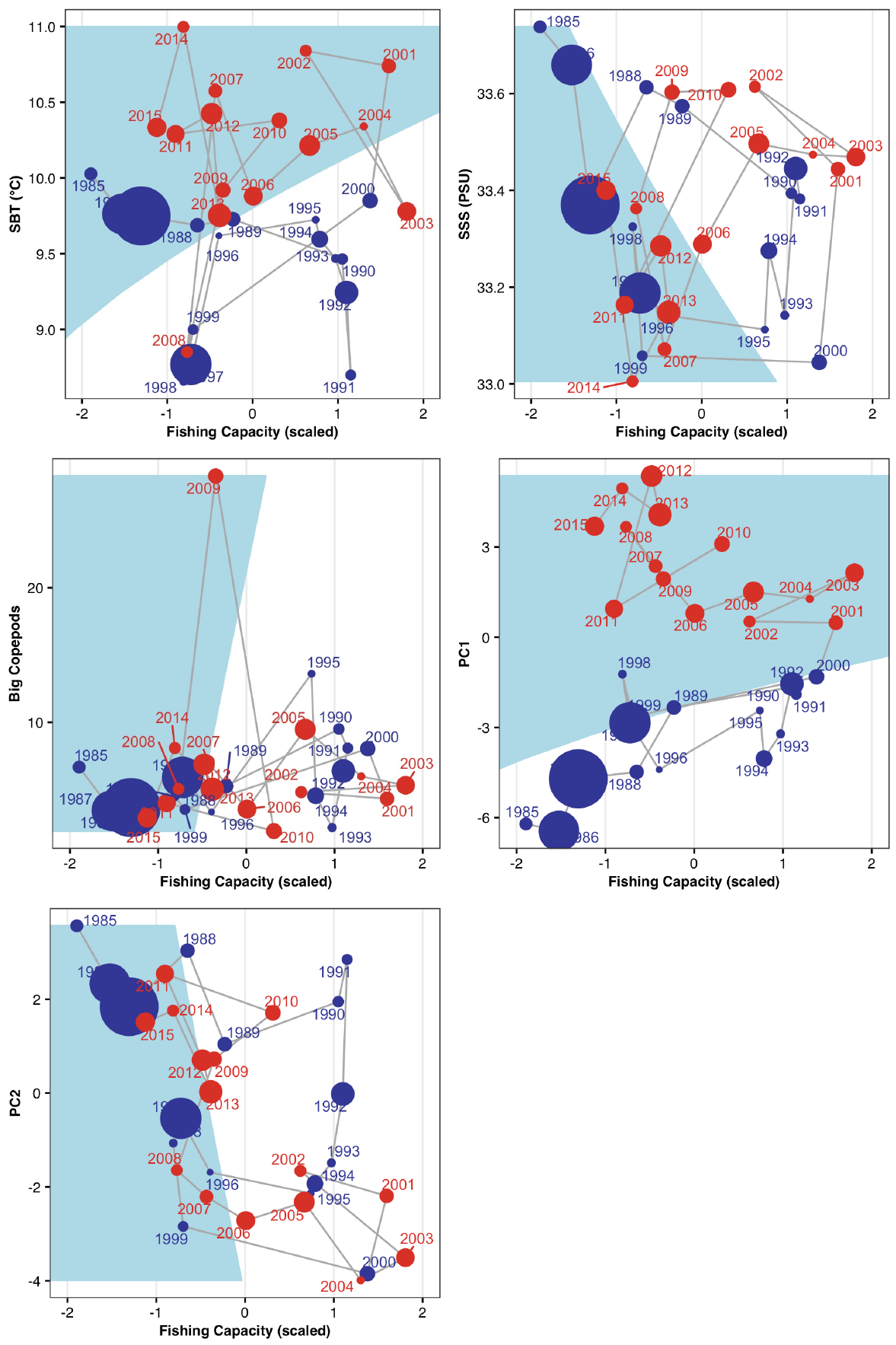

**Figure 9: The stochastic cusp models of the Southern North Sea cod.** Output of the stochastic cusp models with the different beta variables. On the x axis the asymmetry variable, fishing capacity expressed in hours and scaled. On the y axis the bifurcation factors as linear models of the different biotic and abiotic variables. The blue area corresponds to the transition area, the area where three equilibria of the systems are present; two stable and one unstable. The dimension of the dots is proportionate to the CPUE of the cod population (big dot = high CPUE, small dots = low CPUE). The colors of the points represent the periods (blue) before and (red) after 2000.

**Supporting Information C:**

**Analysis of the Southern and North-Western North Sea aggregated area.**

We aggregated the data of North-West and Southern North Sea since these two populations are considered as being part of one population. The time series of the CPUE of cod in this area show a strong decline of cod, especially after 2000 and no sign of recovery (SI Fig.10). The trends of the drivers show a clear increase in temperature and salinity, an oscillatory trend for copepods and a decline of fishing pressure since after the 2000s (SI Fig.11). We performed a PCA to explore how the community of this area changed over time. Recently flatfish, but also rays and other gadoids as hake, became more abundant (SI Fig.12, 13). The driver-state plot shows a strong sign of hysteresis (SI Fig.14). The results of the cusp models indicate contrasting dynamics (SI Tab. 4). After analyzing those results we believe it is more sensible to consider the Southern and North-West area separated. In fact, the dynamics of these two areas are opposite, starting from the time series for the CPUE of Cod (Figure 1 of the main text) and as also showed by the stochastic cusp models, which indicated linear dynamics for the North-West and discontinuous dynamics for the South. Thus, the emergent dynamics of these two different areas are masked when they are combined together, and, in general, the results of the North-Western area are hidden by the Southern area.

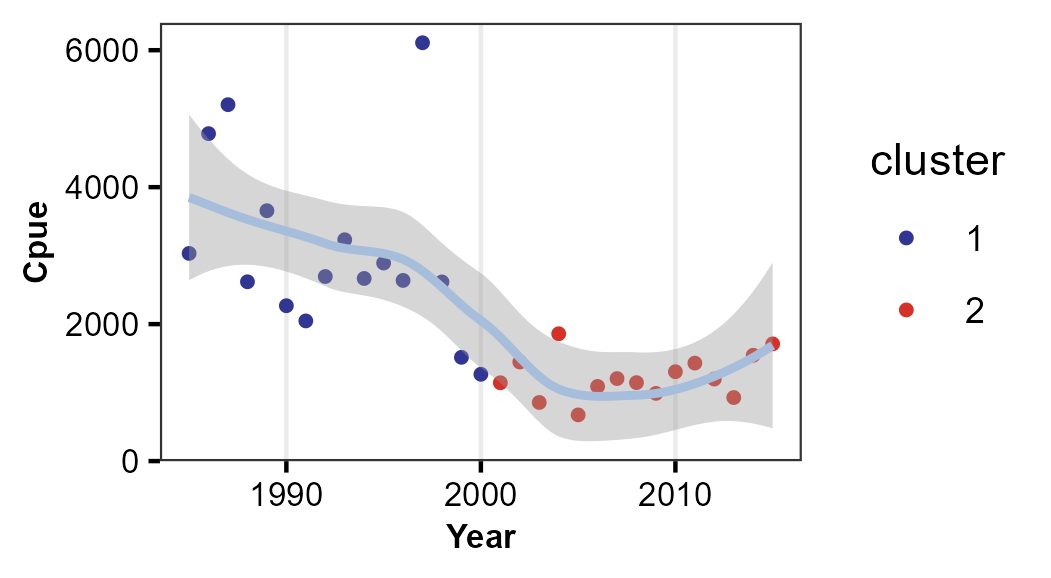

**Figure 10: Time series of CPUE of cod in the aggregated area of the Southern North Sea and the North-West North Sea.** The different colors of the point correspond to the data before 2000 (blue) and after 2000 (red). The line passing through the dots is the smoothed trend done by fitting a loess to better visualize the trend of the populations. In grey is the confidence interval of the line.

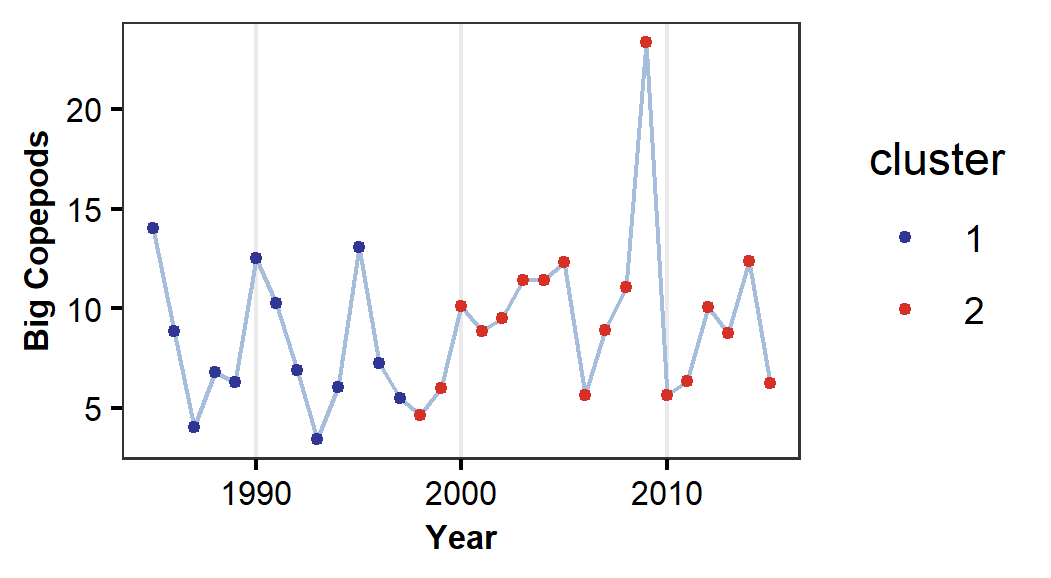

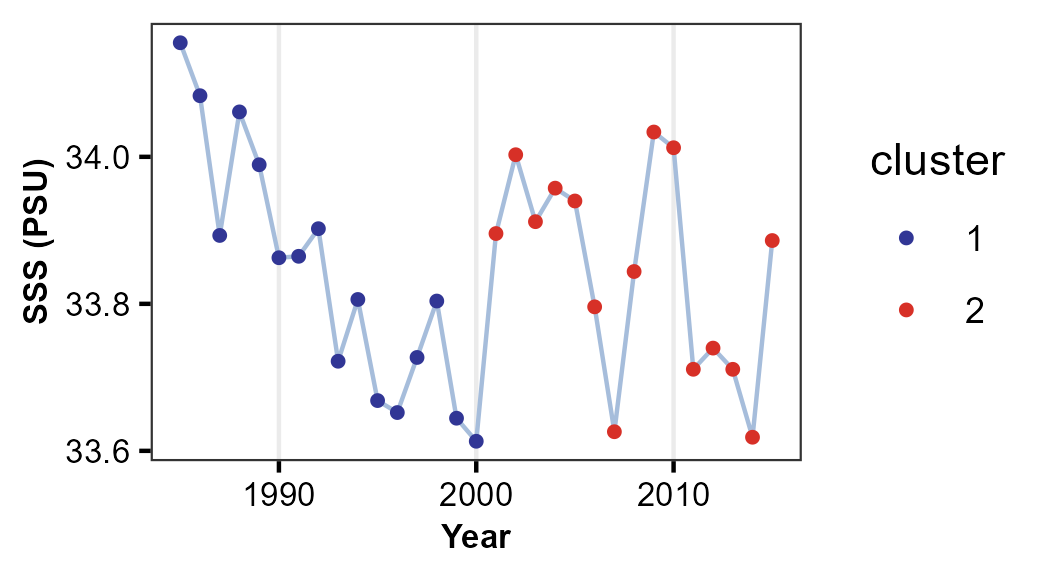

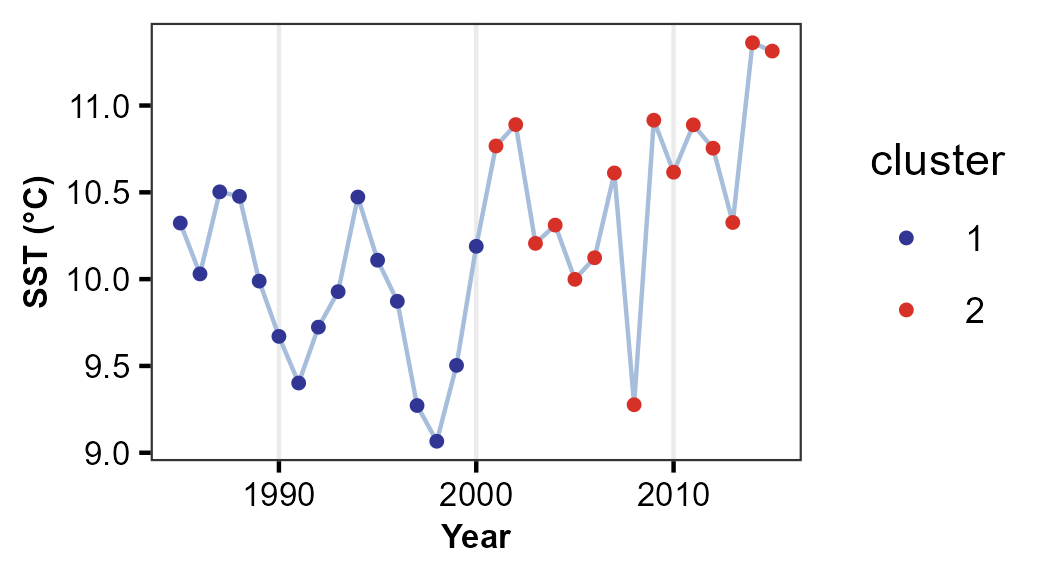

Sea Surface Temperature (°C)

Sea Surface Salinity (PSU)

Big copepods

Effort (thousands of hours)

**a)**

**b)**

**c)**

**d)**

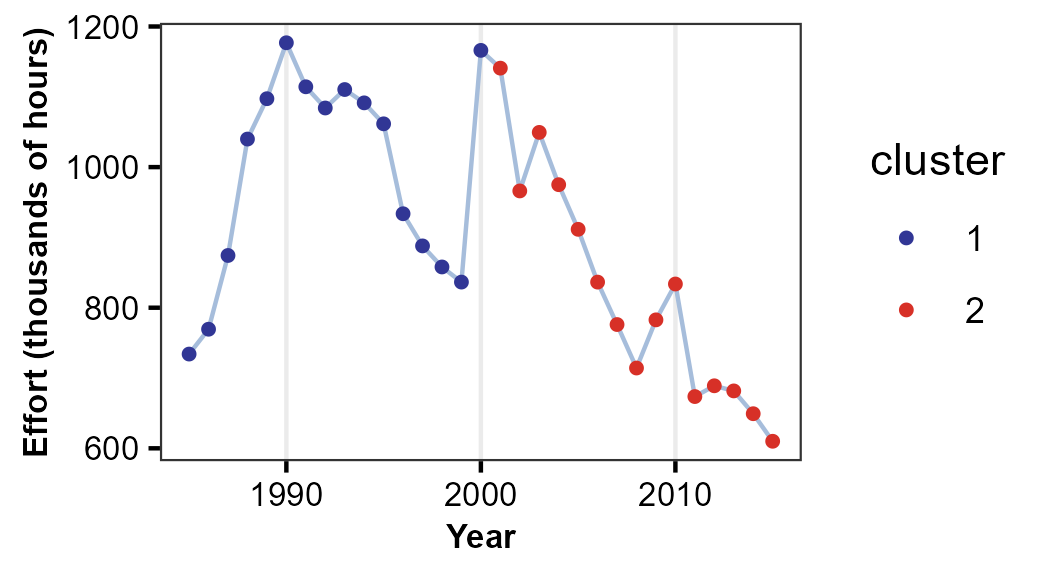

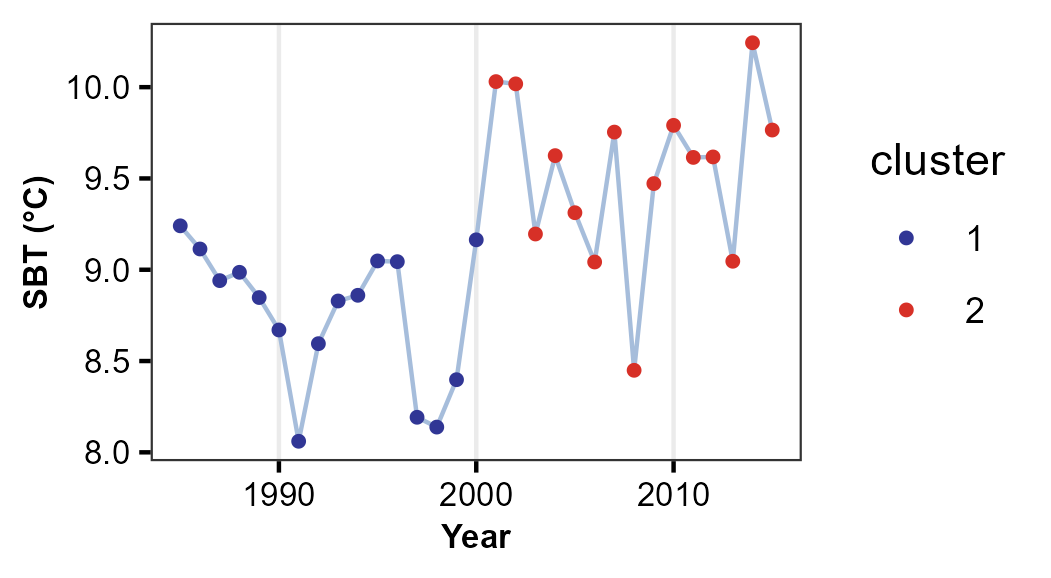

Sea Bottom Temperature (°C)

**e)**

**Figure 11: Anthropogenic, biotic and abiotic drivers of cod in** **the aggregated area of the Southern North Sea and the North-West North Sea.** Time series of effort (a), big copepods (b), Sea Surface Temperature (c), Sea Bottom Temperature (d) and Sea Surface Salinity (e). The color of the dots represents the period before the 2000 (blue) and the periods after the 2000 (red), year in which there were change points.

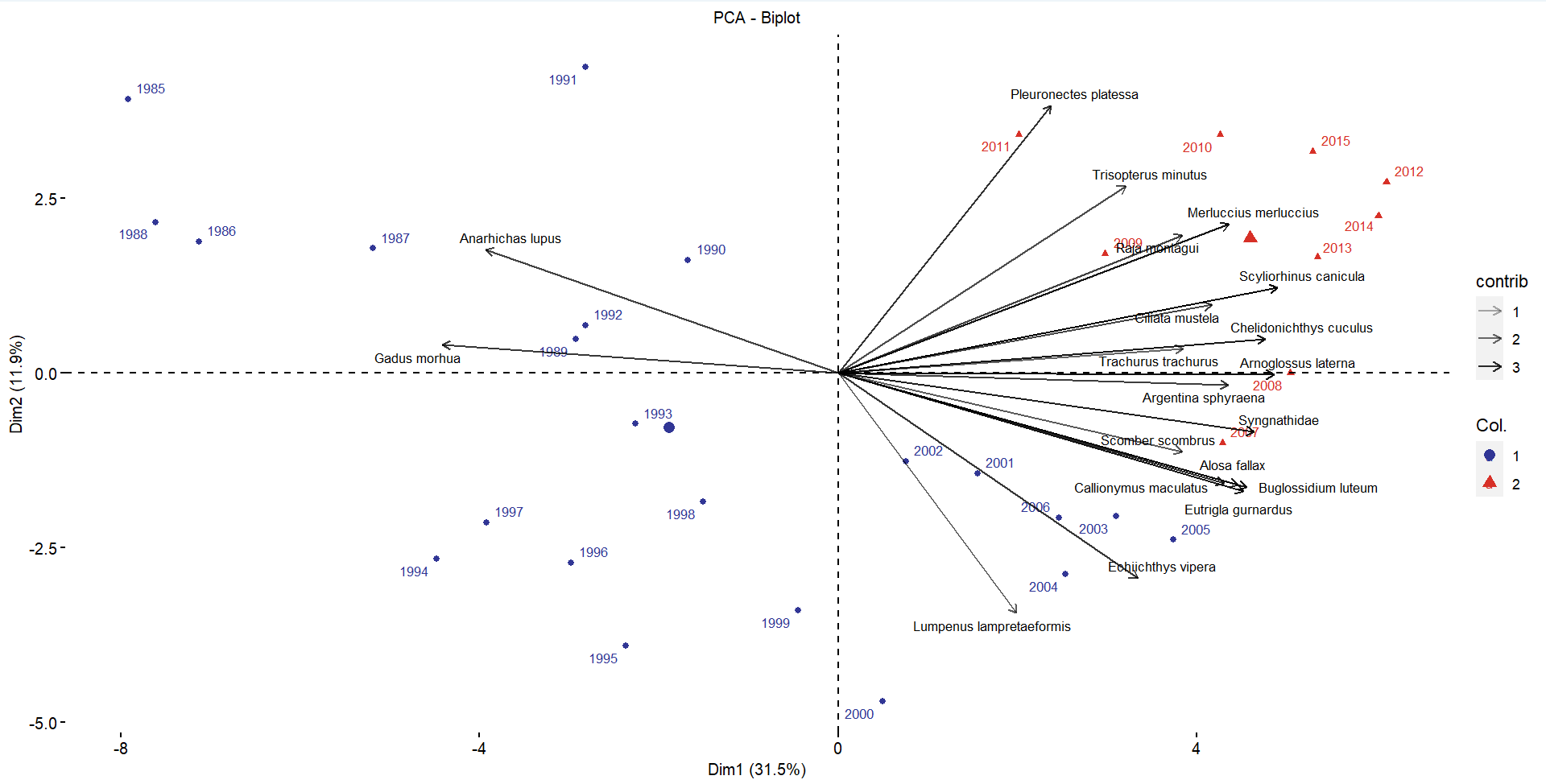

**Figure 12: PCA of the aggregated area of the Southern North Sea and the North-West North Sea**. PC1 and PC2 explain respectively 31.5% and 11.9% of the variance. The years are colored by the clusters identified by the hierarchical clustering analysis (in blue before 2007).The scientific name of the species are shown.

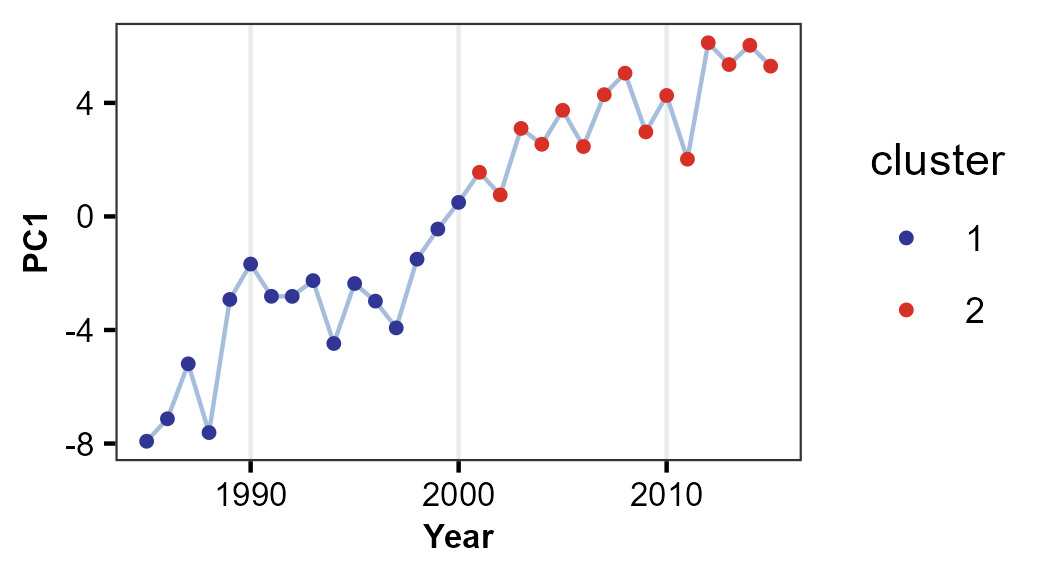

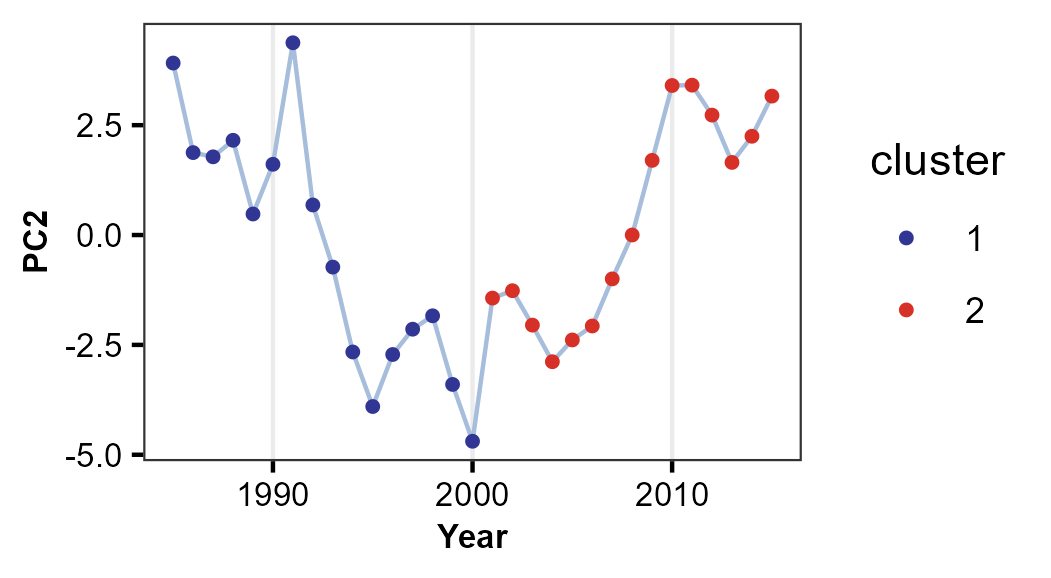

**South-NW aggregated area**

**a)**

**b)**

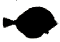

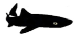

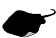

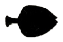

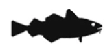

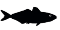

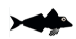

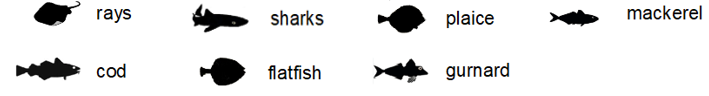

**Figure 13: Community changes in the aggregated area of the Southern North Sea and the North-West North Sea.** Changes in the community as highlighted by the Principal Component Analysis. PC1 (a) and PC2 (b) are used as proxy of the community status. The color of the dots represents the periods before (blue) and after (red) the change point in 2000. The icons show the dominant species for each of the two periods in every area.

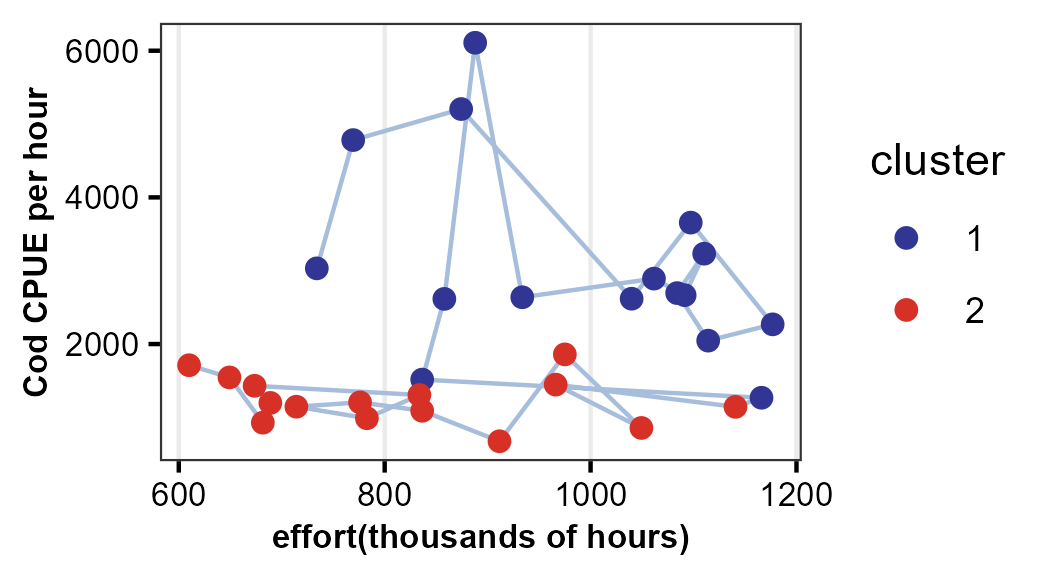

**Figure 14: Hysteresis patterns of the aggregated South-NW cod population.** The driver-state plot using as dependent variable CPUE of cod and as explanatory variable fishing effort (thousands of hours). The different colors of the point correspond to the data before 2000 (blue) and after 2000 (red).

**Table 4. Results of the cusp models for the aggregated South and North-West North Sea data.** Estimates of the parameters of the stochastic cusp model. α1 and α2 are respectively the intercept and slope of the splitting parameter depending on fishing. β1 and β2 are respectively the intercept and slope of the bifurcation parameter that changes in all the models. w1 and w2 are the intercept and the slope of the state variable, the abundance of cod. The significant levels of the estimates are indicated with a star (* p < 0.05, ** p< 0.005, *** p< 0.0005). The R^2^ are pseudo Cobb R^2^ computed to compare the model using the cusp formulation or a linear or a logistic model.

| **Parameters** | **SST** | **SSS** | **SBT** | **Big copepods** | **PC1** | **PC2** |
| --- | --- | --- | --- | --- | --- | --- |
| **α1** | -0.9** | -0.87** | -0.94** | -0.93** | -0.97* | -0.89** |
| **α2** | 0.11 | 0.16 | 0.08 | 0.16 | -0.26 | 0.23 |
| **β1** | -4.7 | 57 | -8.1 | 0.82 | 3.72*** | 1.78** |
| **β2** | 0.64 | -1.63 | 1.11* | 0.12 | 0.57*** | -0.01 |
| **w1** | -1.12*** | -1.06*** | -1.2*** | -1.15*** | -1.74*** | -1.05*** |
| **w2** | 0.9*** | 0.92*** | 0.9*** | 0.9*** | 1.02*** | 0.92*** |
| **R^2^ linear** | 0.11 | 0.03 | 0.19 | 0.15 | 0.6 | 0.36 |
| **R^2^ logistic** | 0.14 | 0.29 | 0.28 | 0.38 | 1 | 0.19 |
| **R^2^ cusp** | 0.27 | 0.35 | 0.16 | 0.13 | 0.55 | 0.34 |
